# RIDGE, a tool tailored to detect gene flow barriers across species pairs

**DOI:** 10.1101/2023.09.16.558049

**Authors:** Ewen Burban, Maud I. Tenaillon, Sylvain Glémin

**Author notes:** Equal contribution (Maud I. Tenaillon, Sylvain Glémin).

## Abstract

Characterizing the processes underlying reproductive isolation between diverging lineages is central to understanding speciation. Here, we present RIDGE – Reproductive Isolation Detection using Genomic polymorphisms – a tool tailored for quantifying gene flow barrier proportions and identifying the corresponding genomic regions. RIDGE relies on an Approximate Bayesian Computation with a model-averaging approach to accommodate diverse scenarios of lineage divergence. It captures heterogeneity in effective migration rate along the genome while accounting for variation in linked selection and recombination. The barrier detection test relies on numerous summary statistics to compute a Bayes factor, offering a robust statistical framework that facilitates cross-species comparisons. Simulations revealed that RIDGE is particularly efficient both at capturing signals of ongoing migration and at identifying barrier loci, including for recent divergence times (~0.1 2*N_e_* generations). Applying RIDGE to four published crow datasets, we validated our tool by identifying a well-known large genomic region associated with mate choice patterns. We identified additional barrier loci between species pairs, which have shown, on the one hand, that depending on the biological, demographic, and selection contexts, different combinations of summary statistics are informative for the detection of signals. On the other hand, these analyses also highlight the value of our newly developed outlier statistics in challenging detection conditions.

## Introduction

The process of speciation involves a gradual and divergent evolution of populations, passing through conditions of semi-isolated species, coined the “grey zone of speciation” (Roux et al. 2016), until complete genetic isolation is achieved resulting in the formation of distinct species (Wu, 2001). Population divergence can occur through various scenarios, ranging from the complete absence of genetic exchanges, known as allopatric speciation (e.g., due to geographical barriers between populations), to almost unrestricted genetic exchanges in sympatric speciation. These extreme scenarios are not mutually exclusive, as genetic exchanges can reoccur after a period of allopatric divergence followed by secondary contacts (Schluter, 2001). Regardless of the scenario, the question of how reproductive isolation is established between divergent populations is central to understanding speciation. This involves comparing the proportion and identity of the relevant genomic regions across biological systems (Delmore et al., 2018; Fraïsse et al., 2021; Schluter, 2001)

Extensive exploration of the genomic bases of speciation have been conducted, in particular in the case of ecological speciation where environmental disparities among populations drive both phenotypic divergence and reproductive isolation (Rundle & Nosil, 2005; Schluter, 2000; Shafer & Wolf, 2013). A recurrently observed pattern is that pre-mating reproductive isolation is facilitated by the physical linkage between genes that govern reproductive isolation and those responsible for divergent traits, which can potentially result from adaptation to contrasted environmental conditions. The gradual establishment of linkage disequilibrium between these genes can then lead to the progressive arrest of gene flow during the speciation process (Schluter & Rieseberg, 2022).

For example, in stickleback fish, divergent mate preferences have been mapped to the same set of genomic regions controlling body size, shape, and ecological niche utilization (Bay et al., 2017). Another striking example concerns the genomic determinants of mate selection based on feather color patterns in carrion and hooded crows (Metzler et al., 2021; Poelstra et al., 2014). Specifically, genes encoding feather pigmentation and genes responsible for perceiving color patterns have been identified within the same 1.95 Mb region of chromosome 18. This region displays significant genetic differentiation between carrion and hooded crows. Similarly, in the neotropical butterflies *Heliconius cydno* and *H. melopomene*, assortative mating behavior is associated with a genomic region proximate to *optix*, a crucial locus influencing distinct wing color patterns between these species (Merrill et al., 2019). Note that, inversions can help build linkage disequilibrium by generating large genomic regions of suppressed recombination, maintaining combinations of co-adapted alleles encoding ecologically relevant traits. For example, in three species of wild sunflowers, 37 large non-recombining haplotype blocks (1-100 Mbp in size) contribute to strong pre-zygotic isolation between ecotypes through multiple traits such as adaptation to soil and climatic conditions or flowering characteristics (Todesco et al., 2020).

Another key genetic mechanism involved in speciation is the epistatic interaction between genes that produce deleterious phenotypes in hybridization, also known as Bateson-Dobzenski-Muller Incompatibility (BDMI) (Gavrilets, 2003). Across *Arabidopsis thaliana* strains, epistatic interactions between alleles from two loci located on separate chromosomes, resulted in an autoimmune-like responses in F1 hybrids (Bomblies et al., 2007). In the Swordtail fish species, *Xiphophorus birchmanni* and *X. malinche,* an interaction between two genes generates a malignant melanoma in hybrids associated with strong viability selection (Powell et al., 2020).

As population-wide genomic data increase, genome-scan approaches enable a more systematic search of the genetic factors behind reproductive isolation. One popular approach relies on the search for genomic islands of elevated differentiation compared with the genomic background, typically through *F_ST_* scans (Wolf & Ellegren, 2017). However, it is now widely recognized that processes other than selection against gene flow can generate such islands. For example, selective sweeps and background selection against deleterious alleles both decrease genetic diversity at linked sites especially in low recombination regions (Charlesworth, 1993; Charlesworth & Jensen, 2021; Cruickshank & Hahn, 2014; Kaplan et al., 1989). Because gene flow barriers are more likely to occur in functional regions, they are also more affected by those forms of selection, further complicating the distinction of gene flow reduction (Ravinet et al., 2017). Demography, which affects the entirety of the genome, is also key to account for barrier detection because barrier loci are harder to identify when the time split is recent and/or the migration rate is low (Sakamoto & Innan, 2019). Yet, recent splits of partially isolated taxa are of paramount interest in speciation research as they allow access to the key determinants of reproductive isolation while avoiding the confusion with other differences accumulated since speciation (Tenaillon et al., 2023).

Linked selection (at least some forms of) can be approximated by a local reduction in effective population size (Cruickshank & Hahn, 2014; Ravinet et al., 2017; Sakamoto & Innan, 2019) and several methods have proposed to decouple its effect from the heterogeneity in effective migration rate to detect gene flow barrier on genomic polymorphism patterns (Fraïsse et al., 2021; Laetsch et al., 2023; Sethuraman et al., 2019; Sousa et al., 2013). These methods relax the assumption that all loci share the same demography. Some of them use likelihood methods to directly estimate and decouple the effects of differential introgression and demography across genomic loci (Laetsch et al., 2023; Sethuraman et al., 2019; Sousa et al., 2013). However, they make specific assumptions about demography. For example, gIMbl simulates population divergence with constant migration, (Laetsch et al., 2023). DILS proposes a more flexible approximate Bayesian computation (ABC) approach (Fraisse et al. 2021). It first infers a demographic model while accounting for heterogeneity in effective population size *N_e_* (to mimic linked selection) and heterogeneity in effective migration *m_e_* (to mimic gene flow barriers), as taking genomic heterogeneity into account has been shown to enhance the quality of model inferences (Roux et al., 2014). Second, the method infers the migration model at the locus scale − arrest of migration vs migration similar to the genome-wide level −, conditioned on the chosen model (Fraïsse et al., 2021). Although effective in detecting gene flow barrier, this dependence on the initial model choice limits comparability among species pairs.

Overall, an adequate method to identify potential reproductive isolation barriers would require a cross-species comparative framework that takes genomic heterogeneity into account, while making analysis comparable despite differences in demographic histories. Here, we propose an innovative method to identify gene flow barrier loci satisfying these requirements and that also quantifies the confidence in locus detection. We used an ABC-based model averaging approach that accounts for different modalities of divergence between pairs of populations/taxons. We considered both heterogeneity in *N_e_*along the genome, by modeling the mosaic effect of linked selection as in the DILS program (Fraïsse et al., 2021), and heterogeneity in recombination, by including an option for the user to provide a recombination map. In addition, we relied on a number of classic summary statistics but also incorporated new ones, related to outlier detection, which improved the inferences of barrier loci. Finally, the method provides Bayes factors associated with barrier detection, which facilitate cross-species comparisons.

## Material and Methods

### RIDGE pipeline

RIDGE utilizes ABC based on random forest (RF) to detect barrier loci between two diverging populations in the line of the framework proposed in DILS (Fraïsse et al., 2021). The observed data consist of a set of loci sequenced on several individuals of the two populations. The general principle of RIDGE is as follows: first, we simulate 14 demographic x genomic models to produce a reference table. This table serves to train a RF that generates weights and parameter estimates for each model according to their fit to the target (observed) dataset. Second, we construct a hypermodel where the posterior distribution of each parameter is obtained as the weighted average over the 14 models. Finally, we use this hypermodel to produce datasets for control loci (thereafter non-barrier) and barrier loci that have undergone no gene flow during divergence. This second set of simulated datasets are employed to train a second RF model that subsequently calculates posterior probabilities and associated Bayes factors for categorizing each locus as barrier or non-barrier.

### ABC Summary statistics

ABC inferences rely on summary statistics that are computed either at the locus-level or across loci, i.e. genome-wide distributions of summary statistics and correlations among loci, and either within or between populations. For a given observed dataset, the number of loci used for the building of the hypermodel is set by the user. To reduce computation time for large datasets, a subset of loci can be randomly sampled to represent the whole genome (by default, we used 1000 loci).

For each locus, RIDGE computes the following within population statistics: the number of Single Nucleotide Polymorphisms, SNPs (S), the nucleotide diversity *π* (Nei & Li, 1979), Watterson’s *θ* (Watterson, 1975), as well as Tajima’s *D* (Tajima, 1989). As measures of population differentiation between populations, RIDGE computes *F_ST_* (Bhatia et al., 2013; Hudson et al., 1992), the absolute *(Dxy)* and the net *(Da)* divergence (Nei & Li, 1979), the summary of the joint Site Frequency Spectrum (jSFS) (Wakeley & Hey, 1997) with *ss* (the proportion of shared polymorphisms between populations), *sf* (the proportion of fixed differences between populations), *sxA* and *sxB* (the proportion of exclusive polymorphisms to each population).

Across loci, RIDGE computes the mean, the median and the standard deviation for each summary statistic described above. In addition, RIDGE computes the Pearson correlation coefficient between *Dxy* and *F_ST_* and between *Da* and *F_ST_*. Regarding specific jSFS status, RIDGE determines the number of loci that contains both shared polymorphisms (*ss* > 0) and fixed differences (*sf* > 0) between populations, *ss^+^sf^+^*and following the same rational *ss^+^sf^−^*, *ss^−^sf^+^*, *ss^−^sf*^−^. These statistics are commonly used in ABC, for example in DILS (Fraïsse et al., 2021). To obtain better insights into the proportion of barriers, we introduced new statistics: the proportion of outlier loci, defined as the proportion of loci that exceeds certain thresholds for *F_ST_*, *Dxy*, *sf* and *Da*, or falling below certain thresholds for *π* and *θ*. The thresholds are determined using Tukey’s fences: *t_min_*=*Q_min_*−1.5∗(*Q_max_*−*Q_min_*) and *t_max_*=*Q_max_* +1.5∗(*Q_max_*−*Q_min_*), for the lower and upper thresholds respectively, where *Q_min_* is the lowest and *Q_max_* the highest quartiles (Tukey, 1977). All summary statistics were computed using the *scikit-alle*l (Miles et al., 2021) and *numpy* (Harris et al., 2020) python packages.

### Coalescence simulations

We simulated the evolution of neutral loci (1000 by default) under 14 demographic x genomic models using the *scrm* simulator (Staab et al., 2015), an efficient *ms*-like program (Hudson, 2002). We stored corresponding simulation parameters as well as all summary statistics in a reference table.

#### Demographic models

RIDGE simulates the split of a single ancestral population of effective size *N_a_*, in two daughter populations of size *N*_1_ and *N*_2_ at time *T_split_*. Four different demographic models are considered as in DILS (Fraïsse et al., 2021) (Figure 1: Demographic and genomic models): (1) strict isolation with no migration (SI), (2) isolation with constant migration rate since *T_split_* (IM), (3) secondary contact with no migration after the split until a secondary contact at time *T_SC_* occurs (SC), and (4) ancestral migration with migration occurring initially and ceasing after time *T_AM_* (AM). Migration *M* (expressed in *N.m* units) is assumed to be symmetrical between the two populations.

**Figure 1:**
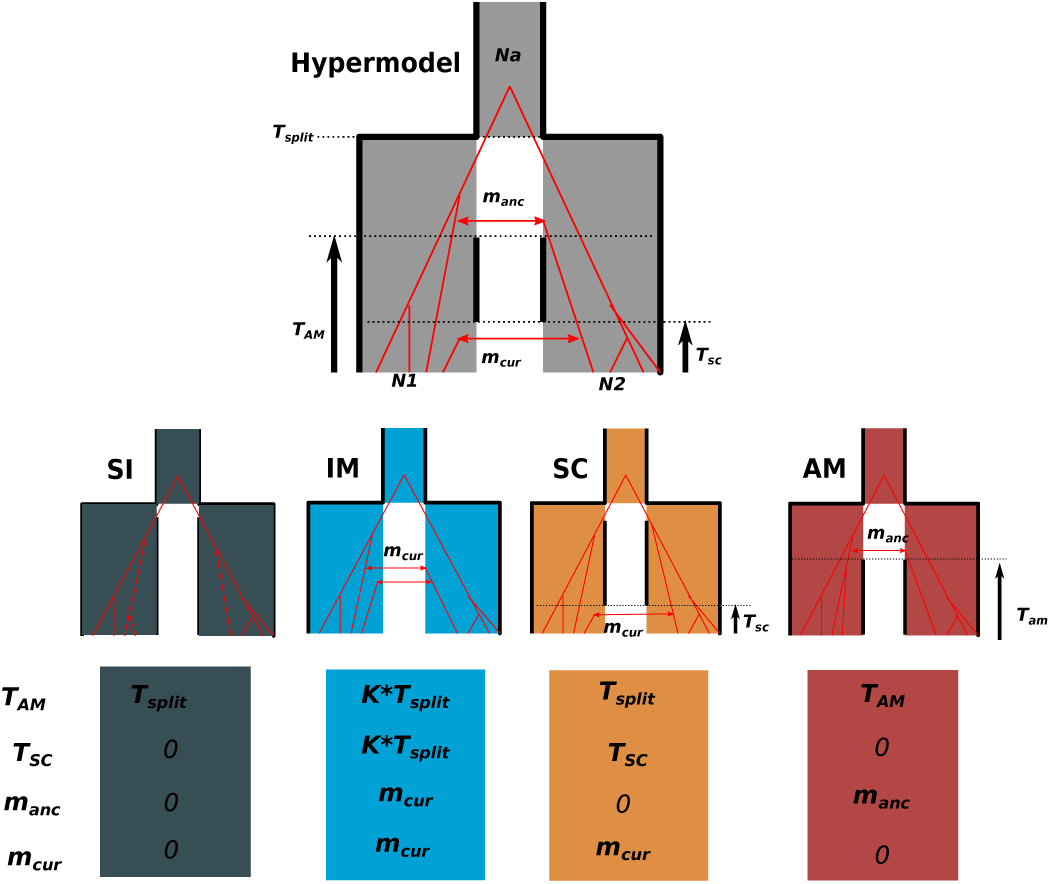
Demographic models implemented in RIDGE. The hypermodel combines all four demographic models considered: Strict Isolation (SI), Ancestral Migration (AM), Secondary contacts (SC) and Isolation-Migration (IM) plus genomic models. In the hypermodel, an ancestral population of effective size Na split at T_split_ in two populations of effective size N_1_ and N_2_. At T_AM_ ancestral migration ceases, and it restarts at the time of secondary contact, T_SC_. M_anc_ and M_cur_ denote the ancestral and current migration rates between populations, respectively. To fit in the hypermodel, each of the four demographic models adopt specific values for four of the parameters as indicated below each graph. For example, under SI, T_AM_ is set to T_split_ as there is no ancestral migration, and T_SC_ is set to 0 as there is no secondary contact, and so are M_anc_ and M_cur_. Note that under IM, in order to model uninterrupted gene flow we considered T_AM_ =T_SC_ =K∗T_split_ where K is a random value drawn from a uniform distribution in [0,1].

#### Genomic models

In addition to modeling demography, RIDGE also incorporates heterogeneity in effective population size along the genome generated by linked selection, and heterogeneity in effective migration generated by selection against migrants at barrier loci. Thus, demographic models are combined with two effective population size modalities (homo-*N* vs hetero-*N*) and with two migration rate (*M*) modalities (homo-*M* vs hetero-*M*) − for models with migration. For simplicity, genomic models are named using a combination of *1N* (homo-*N*), *2N* (hetero-*N*), *1M* (homo-*M*), *2M* (hetero-*M*). While in the *1N* modality all loci display the same effective population size genome-wide, heterogeneity of effective population size under *2N*, is modeled by a rescaled beta distribution. Effective size at locus *i* is given by:

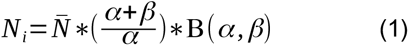

where B(*α,β*) is the Beta distribution with parameter α and β and *N̄* is the mean effective population size across the genome. It is worth noting that for migration (*M*) we fixed the product *N.m* and genome-wide heterogeneity in effective migration is modeled by a Bernouilli distribution where a proportion *Q* of loci displays *M* =0 and a proportion 1−*Q* loci displays *M* > 0, *M* designating either the current migration (*M_cur_*) or the ancestral migration (*M_anc_*). Likewise, we referred to the proportion of barriers under current (*Q_cur_*) and ancestral (*Q_anc_*) migration.

RIDGE assumes that all loci are independent and experience a genome-wide homogeneous mutation rate (*μ*, set by the user) and recombination rate (*r*, set by the user) unless a recombination map is provided, in which case locus-specific recombination rates are given by the recombination map.

### Generation of the reference table

RIDGE explores 14 demographic x genomic models of divergence using a hypermodel that integrates them all. This model contains 12 parameters, eight demographic parameters (*N_a_, N*_1_, *N*_2_, *T_split_, T_AM_, T_SC_, M_cur_, M_anc_*) as described in Figure 1, and four genomic parameters (*α, β,Q_cur_, Q_anc_*). Regarding the demographic parameters, population sizes (*N_a_, N*_1_, *N*_2_) and times (*T_split_, T_AM_, T_SC_*) are sampled in uniform distributions with boundaries specified by the user. Migration rates are drawn from a truncated log-uniform distribution, with the boundary also specified by the user. We used log-normal instead of uniform distributions as migration affects most statistics in a non-linear, multiplicative way. Preliminary simulations showed that it improved the performance of migration estimation. Note that depending on the considered demographic model, some of the parameters are set to 0 (Table S1, Figure 1). For example, under SI, only four demographic parameters are estimated (Table S1). Regarding the genomic parameters, parameters of the beta distribution and the *Q* parameter, are sampled in a uniform distribution where *α, β* ∊[0,10] and *Q_anc_, Q_cur_* ∊[0, *Q_max_*]. *Q_max_*≤1 is the maximal proportion of the genome under gene flow barrier set by the user. RIDGE produces the reference table from a set of simulations with parameters sampled from these prior distributions.

### Point estimates and goodness-of-fit of posteriors

RIDGE utilizes the reference table for training a regression RF model (Raynal et al., 2019). This model produces point estimates for the predicted values of each parameter and assigns weights to simulations based on their proximity to the real data using the *regAbcrf* function. The overall weight for each simulation is calculated as the mean of the weights across all parameters, i.e. joint weights. Subsequently, these joint weights are used to subsample a set of simulations (and their corresponding parameter values) that better match the observed data. This subsample of the reference table is referred to as the posterior table. Note that subsampling of parameters according to the joint weights of simulations effectively accounts for the non-independence of parameters. We evaluated the goodness of fit of the posterior distributions using an enhanced version of the *gfit* function of the *abc* packages (Csilléry et al., 2012), which employs a goodness-of-fit statistics approach described in Lemaire et al (2016) and summarized here. To assess the goodness-of-fit of the posterior *G_post,_* we followed these steps: first, summary statistics (in both observed dataset and posterior table) are normalized by their mean absolute deviation determined from the posteriors table. Then, we computed the Euclidean distance between each summary statistics computed from the observed dataset and those computed from each η simulation contained in the posterior table. Together it form a vector of Euclidean distances *d*_1_… *d_η_* on which we computed the average, denoted *D_post_*. To derive the null distribution of *G_post,_* we considered a dataset randomly sampled in the posterior table as “observed” and discarded from subsequent analyzes. The remaining *η* – 1 datasets of the reference table were used to compute D_post_’, the average euclidean distance between the posterior table and the “observed” dataset. Repeated as such *Z* times, we obtained a vector of 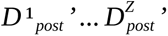. Then, we computed *G_post_* as the proportion of values for which *D_post_’* > *D_post_*.

### Detection of barrier loci

Each set of parameters of the posterior table is used to generate two sets of individual-locus simulations, one set for non-barrier loci (*M* equals to the value of the posterior table) and one set for barrier loci (*M* set to 0), with two corresponding per-locus reference tables. The RF algorithm (*abcrf* package) was trained on these per-locus reference tables to predict the most probable status of each locus, either barrier (model *x_1_*) or non-barrier (model *x_2_*). Since there are only two models, the posterior probabilities satisfied: *P*[*x*_1_]=1−*P*[*x*_2_] so that we were able to compute a Bayes Factor (BF) for each locus *i*, denoted as *BF_i_*:

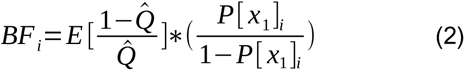

Here, *E*[] represents the average of the ratio (1−*Q̂*)/*Q̂* over the posterior distribution obtained from the hypermodel.

### Evaluation of RIDGE performance on pseudo-observed datasets

We evaluated RIDGE performance on pseudo-observed datasets. As a first step, we evaluated the ability of RIDGE to correctly infer demographic x genomic models. We next used the pseudo-observed datasets to evaluate the accuracy of RIDGE in estimating the proportion of barrier loci, and detecting their locations throughout the genome.

We simulated pseudo-observed datasets under the four demographic models and under both *2M2N* and *1M2N* genomic models (only *1M2N* for SI). For simplicity, we fixed *N_a_*=*N*_1_ =*N*_2_=50000 individuals. The time of the secondary contact (*T_SC_*) was set to 0.2∗*T_split_* and the time of arrest of ancestral migration (*T_AM_*) was set to 0.7∗*T_split_*. We used a range of parameter values (Table S2) for divergence (from 1000 to 2 million generations, i.e. from 0.1 to 20 in 2*N_e_* generation unit), for migration (*M* = 1 and 10 *N.m*), and barrier loci proportion (*Q* = 1%, 5% or 10%). We set the mutation rate to *μ* = 1.10⁻⁸ and the recombination rate to *r* = 1.10⁻⁷ so that their ratio was 10. In total, we simulated 15,000 datasets using the *scrm* coalescent simulator (Staab et al., 2015). Each multilocus dataset contained 1000 loci of 10kb each, and we performed 100 replicates per scenario. To evaluate the inference of demographic x genomic models, we calculated the goodness-of-fit of the estimated model and determined the contribution of each model to the estimation of posteriors obtained from pseudo-data sets. Contributions were evaluated through four criteria: (i) the average weight of the simulated demographic (among the four) model called here the “correct” model, (ii) the average weight of *2M* models, (iii) the average weight of *2N* models, and (iv) the average weight of models displaying current migration. We also compared the point estimates obtained from simulations with the input parameter values.

Next, we assessed our ability to detect barrier loci using the Area Under the Curve (AUC) of the Receiver Operating Characteristic (ROC) curve. The ROC curve relates the false positive rate (FPR) to the true positive rate (TPR) and provides insights into the discriminant power of a method. The AUC of the ROC ranges from 0 to 1. An AUC of 0.5 indicates that FPR and TPR are equal irrespective of the threshold, which implies a random classification of loci into barrier and non-barrier loci while an AUC of 1 indicates perfect classification. Additionally, we computed the precision as the number of true positives (TP) divided by the sum of true positives and false positives (TP + FP).

### Application to experimental data on crow hybrid zones

To assess the performance of RIDGE on experimental data, we focused on two published datasets produced by Poelstra et al. (2014) and Vijay et al. (2016). All sequencing data from crows were extracted from NCBI database under project number PRJNA192205 and the reference genome used to map them is GCF_000738735.1. In the first one, a comparison was made between 30 individuals of *Corvus corone* (carrion crows) populations from Spain and Germany, and 30 individuals of the *C. cornix* (hooded crows) population from Poland and Sweden. In the second one, three crow contact zones, among which two well-characterized hybrid zones, with similar divergent times around ~ 80 000 generations are described, from the most recently-diverged pair *C. corone* - *C. cornix* (RX), to the most anciently-diverged *C. cornix* - *C. orientalis* (XO) and *C. orientalis* - *C. pectoralis* (OP) pairs (Vijay et al., 2016). This dataset consisted of 124 sequenced individuals. The number of individuals sampled varied for each pair (RX: 15-14 individuals; XO: 6-6 individuals; OP: 5-3 individuals).

All alignments were done on a reference genome (NCBI assembly: GCF_000738735.1) consisting of 1299 scaffolds resulted in the detection of 16,064,921 common SNPs with an average density of 15 SNPs per kilobase. Previous genome-wide scans across the three pairs identified a number of candidate loci potentially involved in population/species divergence (Vijay et al., 2016). Two metrics were employed in those scans: (i) a Z-transformed *F_ST_* computed on 50 kb non-overlapping windows between population/species pairs and normalized by the local level of Z-transformed *F_ST_* from allopatric pairs, denoted as *F_ST_*’, (ii) an unsupervised genome-wide recognition of local relationship pattern using Hidden Markov Model and a Self Organizing Map (HMM-SOM) method implemented in Saguaro (Zamani et al., 2013) to identify local phylogenetic relationships based on matrices of pairwise distance measures, across each of the target hybrid zones.

Here, we applied RIDGE on 50 kb non-overlapping windows considering a mutation rate of 3.10⁻⁹ for both datasets as is Poelstra et al (2014) and Vijay et al (2016). We therefore focused on scaffolds longer than 50 kb, which accounted for 9% of the total scaffolds but represented 98% of the genome. Prior bounds are given in Table S3, and were determined based on the observed datasets and results of analysis from Vijay et al (2016). First, we compared Bayes factor outliers (BF > 50) from RIDGE results with outlier loci detected in (Poelstra et al., 2014) to assess the ability of RIDGE to correctly detect barrier locis. Secondarily, we analyzed RIDGE results produced on three species pairs on a lager dataset (Vijay et al., 2016) to understand how BF correlate with summary statistics and which summary statistics are able to discriminate outlier loci (BF > 50).

## Results

### Demographic inferences

The RIDGE’s ability to infer demographic parameters, measured by the goodness of fit of posteriors (*G_post_*), far exceeded the rejection threshold of 5% and was stable across all models and conditions tested in pseudo-observed datasets (Figure 2 & S1). However, the model’s contribution to the estimation of the demographic and genomic parameters varied across conditions. The percentage of simulations correctly attributed to the correct model increased with the time split (*T_split_*), reaching over 65.7% for IM, 81.1% for SC and 69.5% for AM models when *T_split_* exceeded 10⁶ generations (Figure 3). In contrast, the SI model never achieved more than 32.1% accuracy. The percentage of simulations correctly detecting the presence or absence of current migration increased with *T_split_* and heterogeneous migration was better captured under current rather than ancestral migration (82.1% and 80.8% at 10⁶ generation for IM and SC against 48.7% for AM). Heterogeneity in population size (2*N*) followed the same pattern across *T_split_*, irrespective of the demographic model. These results indicated that while the correct demographic model was accurately inferred only under specific conditions, the occurrence of current migration was generally well captured.

**Figure 2:**
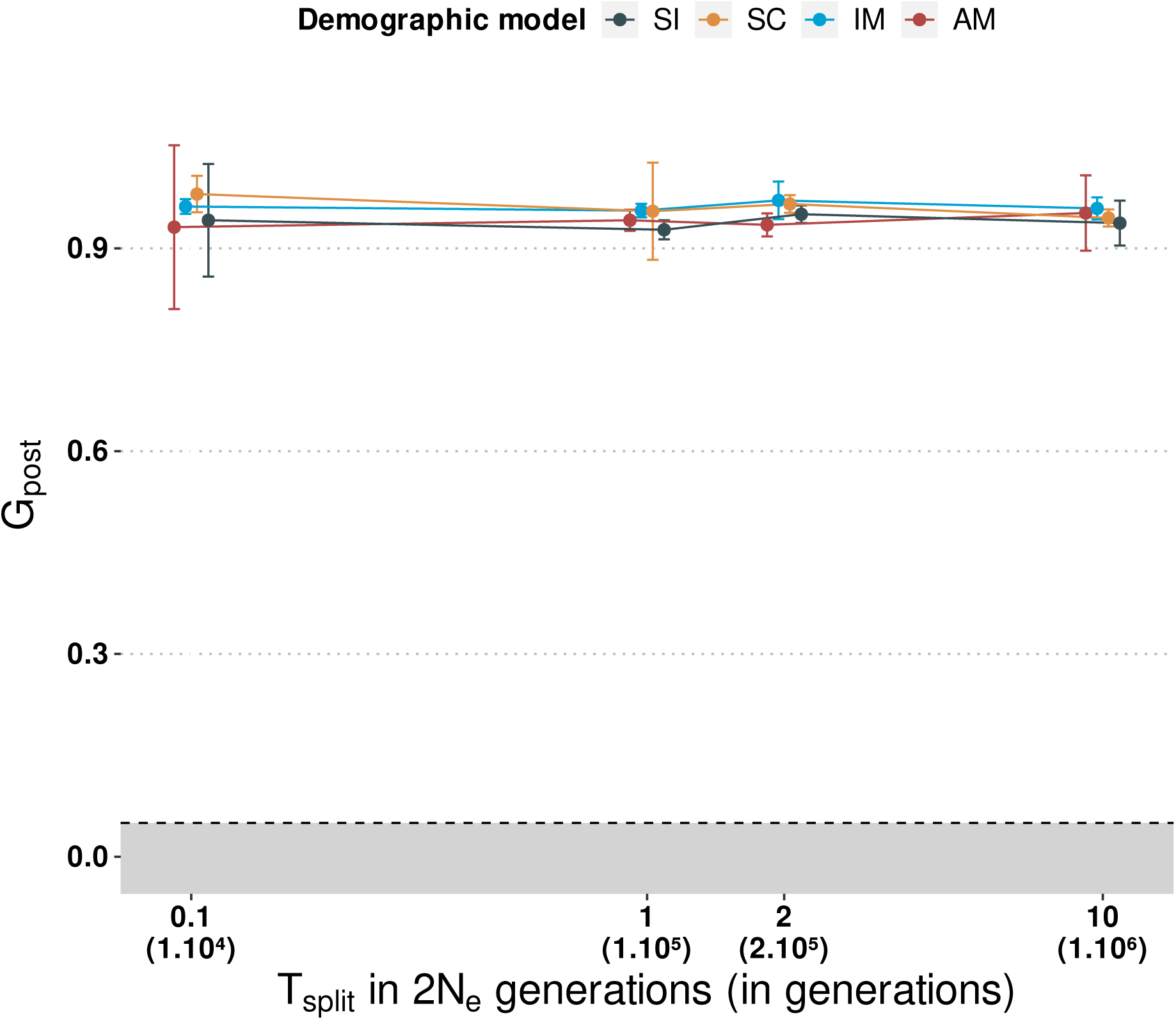
Evolution of the goodness-of-fit of the posteriors (G_post_) as a function of time split, for four demographic models. The rejection threshold of 5% (under which an inferred model is discarded) is represented by the gray zone. Average values over 100 replicates with error bars (standard deviation) are presented. The data used in this figure were obtained from pseudo-observed datasets simulated under the 2N2M model with migration set to 10 (M =10) and a proportion barrier Q=10 % (except for SI, no migration and no barrier).

**Figure 3:**
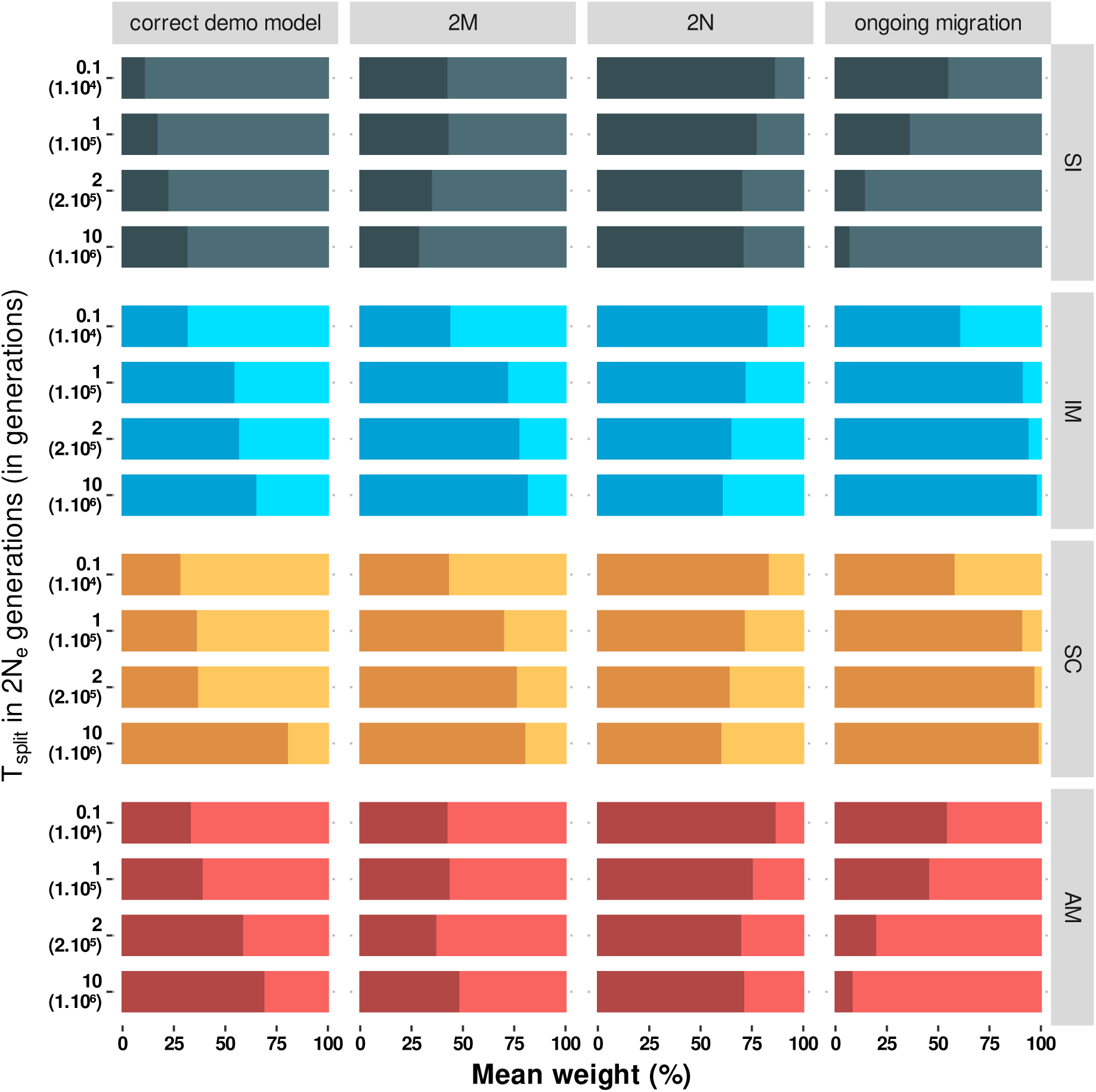
Demographic x genomic model weights in posteriors across time splits. Weight was measured by considering four criteria: i) the average joint weight of the true demographic (among the fours) model –called here the “correct” model– in posteriors, ii) the average joint weight of 2M models, iii) the average weight of 2N models, iv) and the average weight of models displaying ongoing (current) migration. Proportion of accurate model predictions are shown in dark colors. As an example, for a time split of 10⁶, an average weight of 0 for ongoing migration under the SI model signifies that across 100 replicates, simulations under ongoing migration represent 0% of the posteriors and so did not contribute to parameter estimation. All models were simulated under 2N2M, and M_curr_ or M_anc_=10

We also examined the specific point estimates associated with each parameter. The accuracy of *T̂_split_* estimation was only slightly affected by the proportion of barriers and migration rate, closely approximating the simulated value irrespective of the demographic model (Figure S2). Similar patterns were observed for *T̂_SC_* and *T̂_AM_* (Figure S3). As *T_split_* increased, estimates of current population sizes *N̂*_1_ and *N̂*_2_ improved, approaching simulated values when *T_split_* reached 1.10⁵ generations (Figure S4). Estimates of past population size *N̂_A_* is theoretically possible if *T_MRCA_* ≈4 *N_e_* (with *T_MRCA_* the coalescent time of the Most Recent Common Ancestor), if not, all individuals coalesce before *T_split_* so that no signal is available for *N̂_A_*. In our case, *T_MRCA_* ≈4 *N_e_*=2.10⁵ generations, and *N̂_A_* deteriorated beyond this value, converging towards the prior mean (Figure S4). Current migration estimates (*M̂_curr_*) were more reliable than ancestral migration ones (*M̂_anc_*). The proportion of barriers had minimal impact on *M̂_curr_*, under SC and IM models. Deeper *T_split_* resulted in greater migration signal and therefore improved the accuracy of *M̂_curr_* (Figure S5 & Figure S7 left). In contrast, *T_split_* had no clear effect on *M̂_anc_* (Figure S6 & Figure S7).

### Inferences of barrier proportion

The barrier proportion estimate, *Q̂*, plays a crucial role in the computation of Bayes factors (Eq 2) and the detection of barrier loci. We obtained reliable estimates of the barrier proportion, *Q̂*, when there was current migration (IM and SC models) and when *T_split_* exceeded 1.10⁵ generations (Figure 4 & S8). For more recent *T_split_* (< 0.2 2*N_e_* generations, approximately), *Q̂* was not properly estimated and converged to the prior mean, indicating that RIDGE lacks power to discriminate between barrier and non-barrier loci. When there was only ancestral migration (AM model), *Q̂*was not reliable whatever the conditions, except for both high migration rate and divergence time. Under the SI model, for which the proportion of barriers has no significance, the estimates corresponded to the prior mean. The *Q* parameter had a minimal impact on the total effective migration rate, as shown in Figure S7 and S8, and was therefore expected to exhibit a weak correlation with the genome-wide level of genetic differentiation/divergence between populations, as measured by statistics such as *F_ST_*, *Da*, and *Dxy*. We therefore introduced additional summary statistics based on the proportions of outliers for *F_ST_*, *Da*, *Dxy*, *sf* and *π*. To assess the usefulness of these new statistics, we compared *Q̂*estimated with or without them. Overall, outlier statistics reduced estimation errors by 11%. They were particularly effective in improving *Q̂* under challenging conditions for barrier proportion estimation, such as when migration was low (*M* ⩽1) and the proportion of barriers was small *Q*⩽1 % (Figure S9). The impact of outlier statistics varied across models and *T_split_* values. Under the AM model, *Da* outliers positively correlated with *Q̂*(pearson r > 0.56), while under the IM model *sf* outliers exhibited a positive correlation with *Q̂* (*r* > 0.77). For the SC model, correlations between outliers statistics and *Q̂* highly depend on divergence time (Table S4). For the recent time split, there was a positive correlation with *Dxy* (0.68), for intermediate time splits there was a correlation with *Da* (>0.97), and for the oldest time split, there were positive correlations with *F_ST_*, *Da*, and *π* (> 0.99, Table S4).

**Figure 4:**
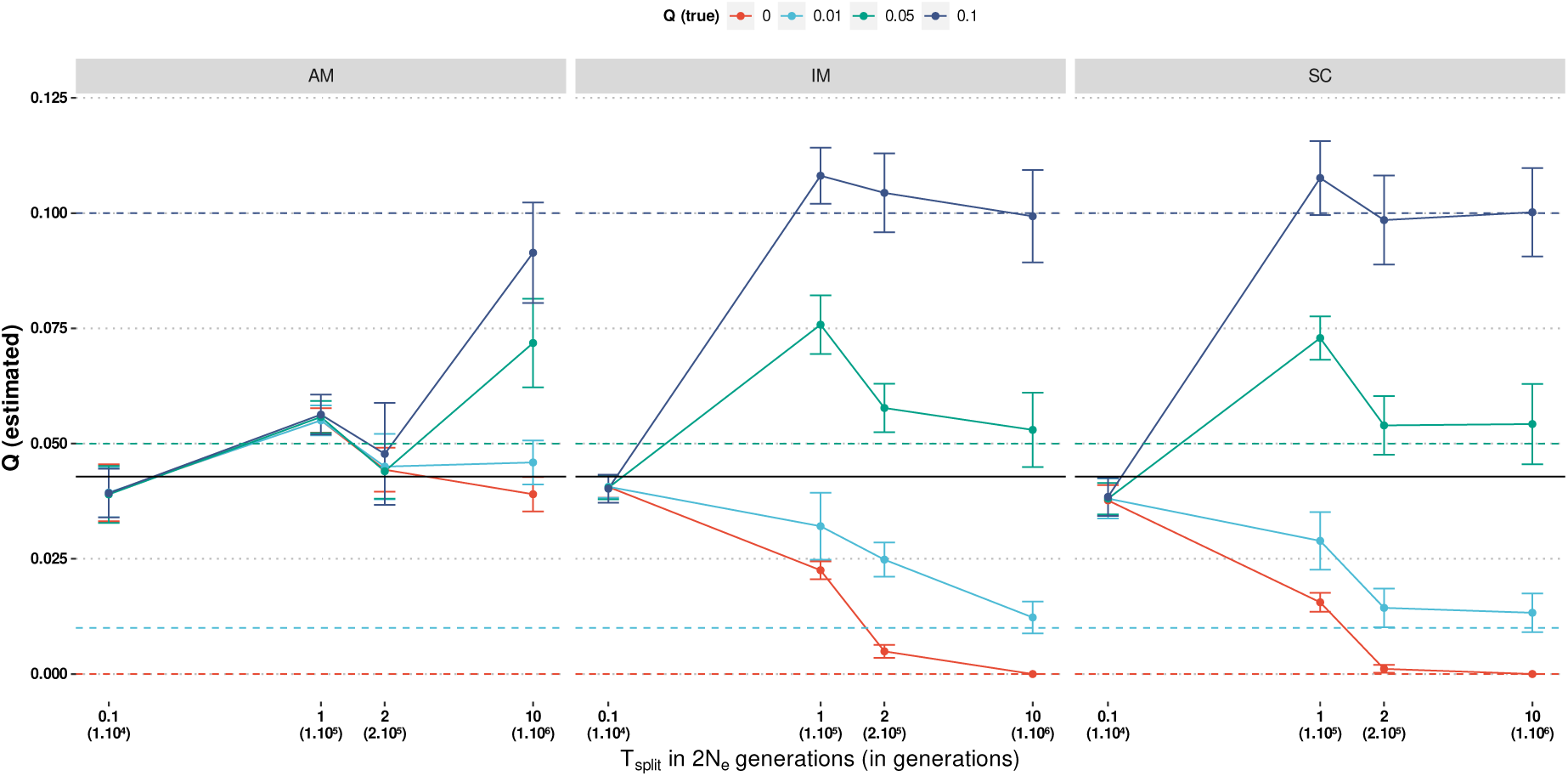
Barrier proportion estimates as a function of divergence time under three demographic models. In this figure, migration is set to M =10 and the plain black line represents the priors mean. Each data point represents the average value over 100 replicates with standard deviation as error bars. Results overall conditions explored are represented in Figure S8.

### Detection of barrier loci

The parameter *T_split_* plays a crucial role in detecting gene flow barriers. This is because the contrast between gene flow barriers and the rest of the genome increases with *T_split_* as illustrated in Figure 5A. As *T_split_* increased, the overlap between the space of summary statistics occupied by barrier and non-barrier loci decreased and correlated with the between corresponding BF distribution (Figure 5A & B). To quantify the discriminant power of RIDGE, we used the area under the curve (AUC) of the receiver operating characteristic (ROC), as depicted in Figure 5C. When *T_split_* was low, the AUC remained close to 0.5, indicating no power to detect barriers. Our results on pseudo-observed data demonstrated that both the ability to detect barriers (measured by the AUC of the ROC) and the precision in barrier detection (measured by the PV/P ratio) increased with *T_split_* (Figure 6). Moreover, barriers were more efficiently detected and at lower *T_split_* under current (IM and SC models) than ancestral gene flow (AM model) as shown in Figure S10 & S11. Noteworthy, in some instances, RIDGE failed to detect any barrier (e.g., when *T_split_*=1.10^4^), in agreement with AUC close to 0.5 (Figure S10). Nevertheless the AUC never dropped below 0.5, indicating that RIDGE did not generate an excess of false positives (Figure S10 & S11).

**Figure 5:**
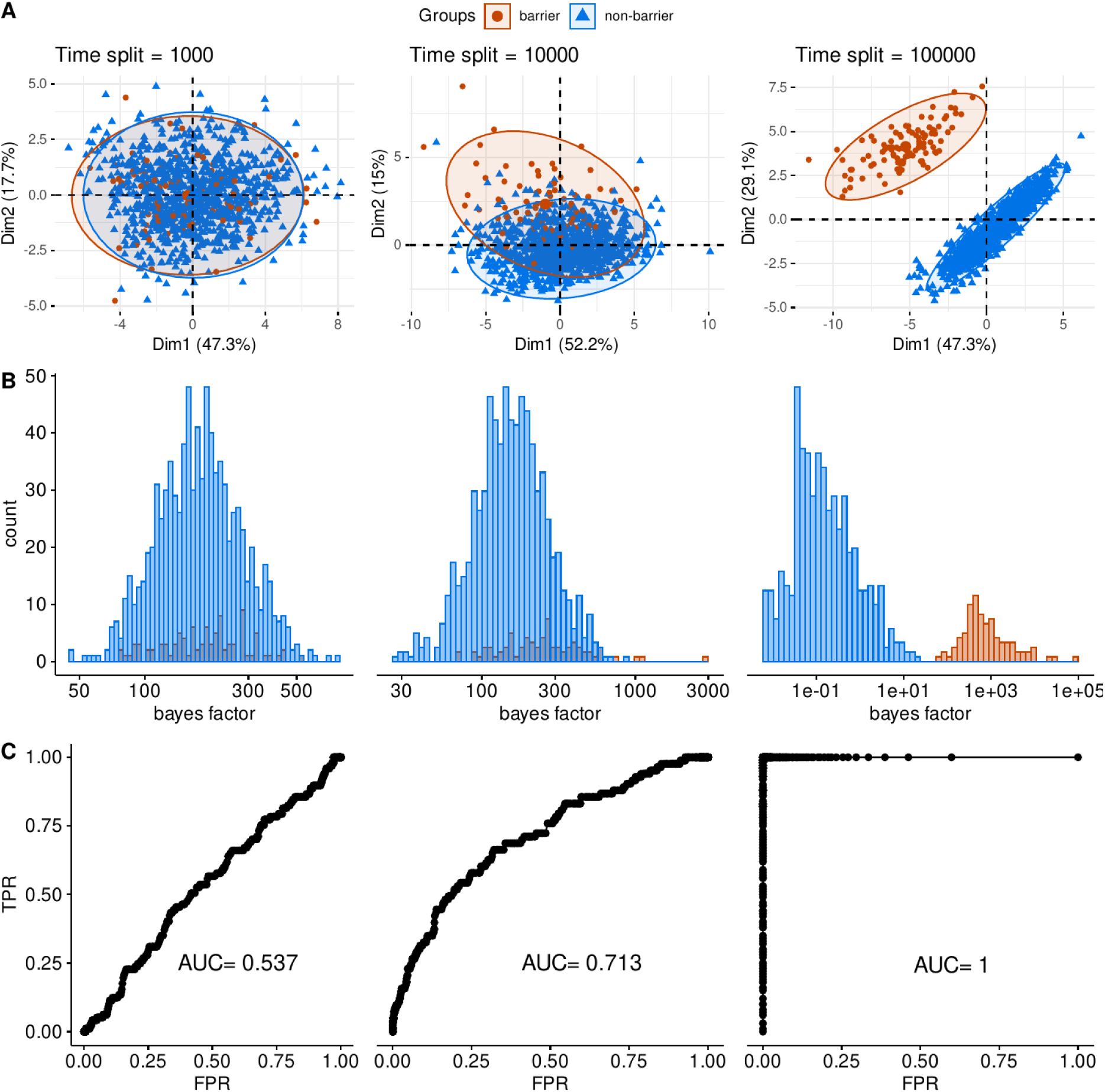
Impact of the divergence time on the overlap between barrier and non-barrier loci. Overlap revealed by a principal component analysis (PCA) computed on all 14 summary statistics (A), the log of the bayes factor (BF) produced by RIDGE (B) and the area under the ROC curve (AUC) of the bayes factor (C). The greater the AUC the higher the discriminant power is. A single pseudo-observed dataset was used for each of the three values of T_split_. Datasets were simulated under an IM 2M2N model, with the following parameters: M =10, and Q=0.1.

**Figure 6:**
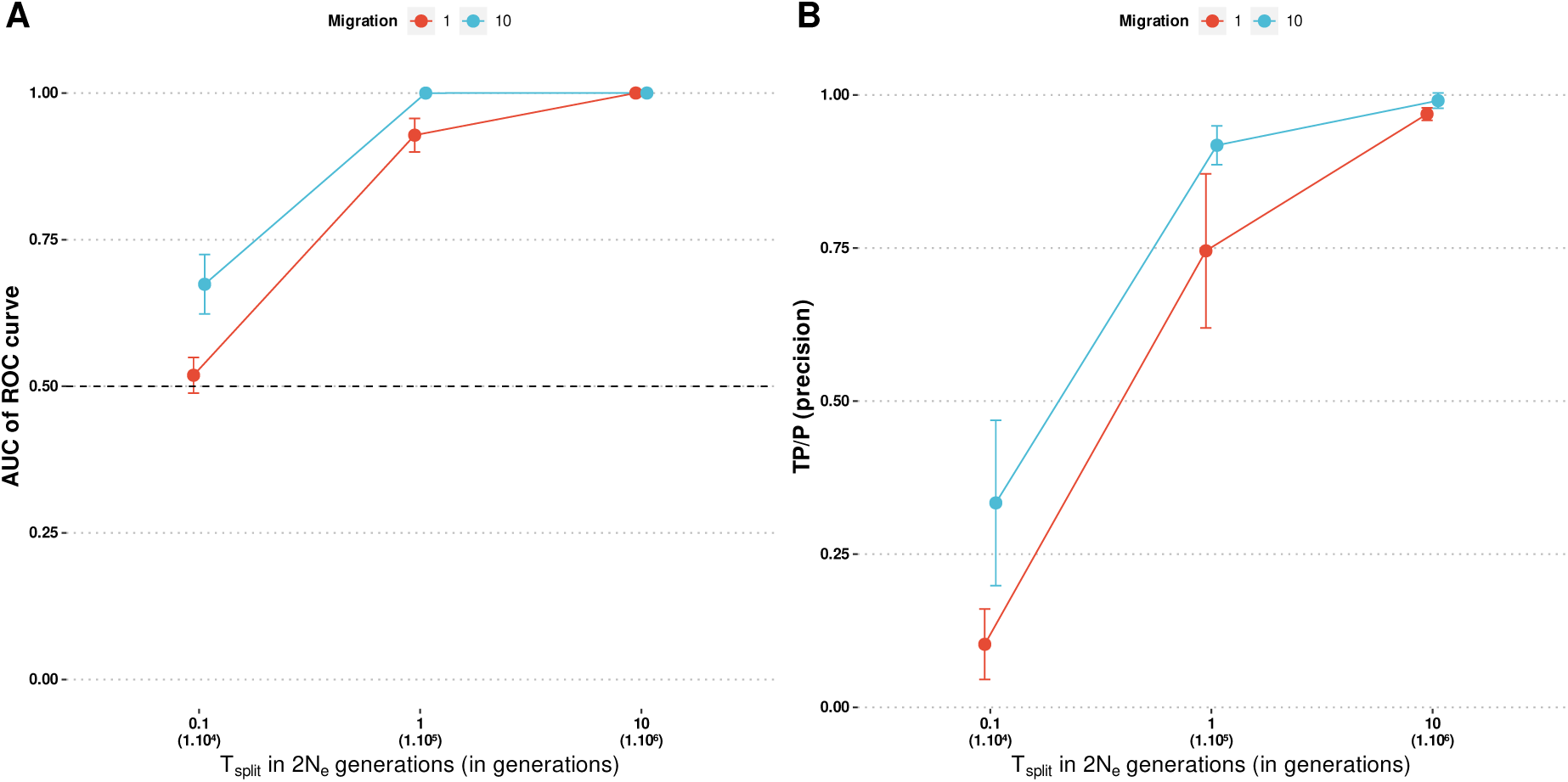
Ability and precision in the detection of barrier loci as a function of divergence time and migration. Ability is measured by the AUC of the ROC (A) and precision by TP/P (B). Considering a proportion of barrier Q̂, barrier loci are those displaying a Bayes factor superior to the quantile at 1−Q̂. Each data point represents the average value over 100 replicates with standard deviation as error bars. Simulations were performed under an IM 2M2N modelwith Q=0.1

### Detection of barrier loci on crows datasets

Poelstra et al (2014) identified a highly divergent region on scaffolds 78 and 60, which contained multiple genes identified through genomic scan, functional analysis, and differential expression. These genes are involved in the melanogenesis pathway and visual perception. This region was thus considered by the author as a “speciation island” allowing for the maintenance of phenotypic differences between crows based on color phenotypes and color-assortative mate choice.

We ran RIDGE on the same dataset using the same window size as in Poelstra et al (2014). Our analysis successfully fitted the observed data, with a goodness of fit indicated by *G_post_* = 0.67. The estimated value of *T̂_split_* in 2*N_e_* generation is *T̂_split_* / 2 *N̂_e_*=0.48, indicating that we were within a favorable range for RIDGE to effectively detect gene flow barriers. The distribution of Bayes Factors (BF) was clearly bimodal with a distinct group of outliers (*BF* >50), which accounted for 0.3% of the genome (Figure 7B). Interestingly, among these outlier loci, four genes (CACNG1, CACNG4, PRKCA, and RSG9) were also found by Poelstra et al (2014) and located on scaffold 78 (Figure 7C). The probability of detecting the same four genes just by chance was low (*p* = 3.59 10^−5^).

**Figure 7:**
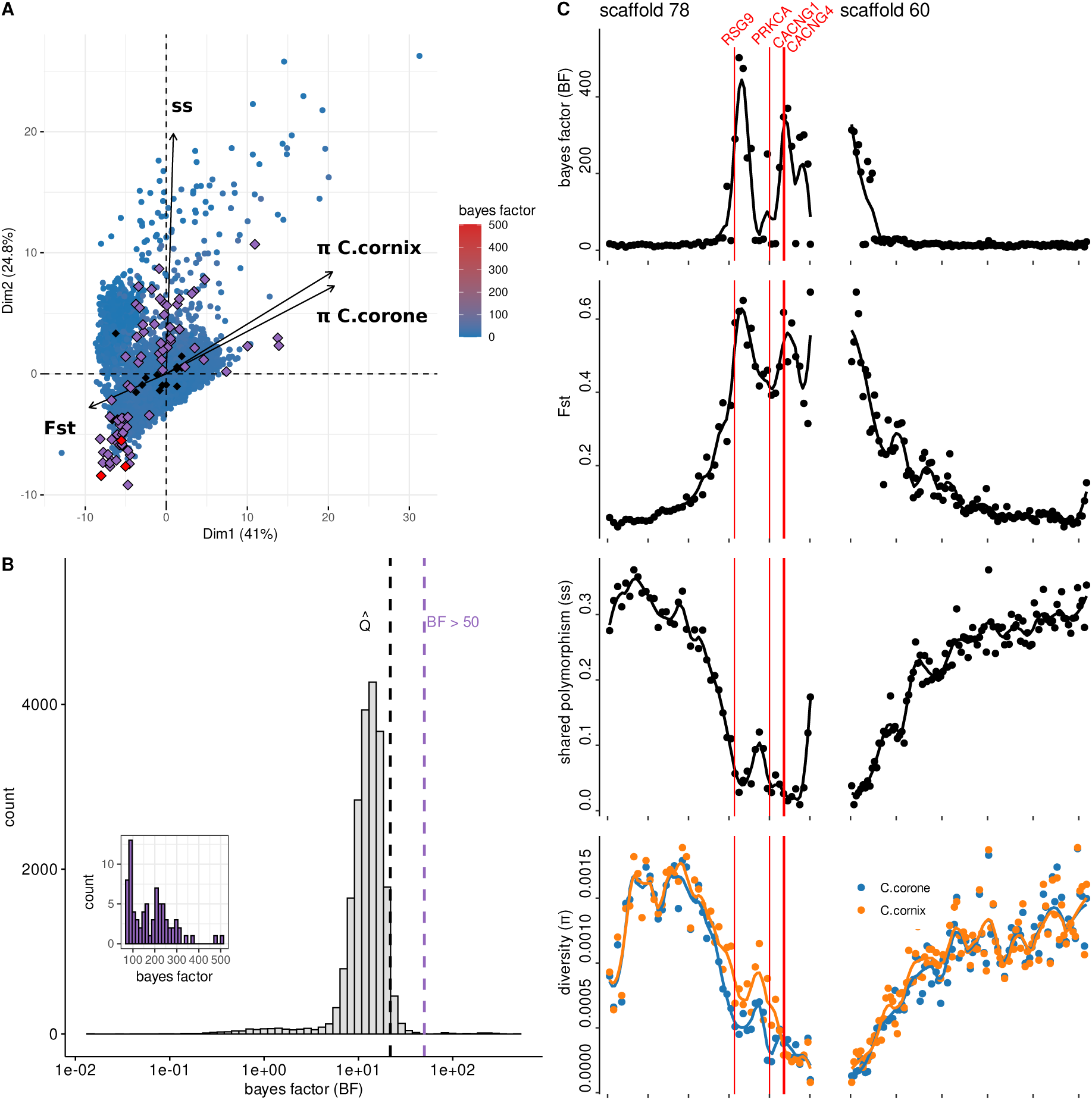
Results of the analysis conducted using RIDGE on the crow hybrid zone between carrion and hooded crows. PCA plot of the summary statistics (only 4 of 14 summary statistics are represented), where each point represents a locus and is color-coded based on its corresponding Bayes factor value (A). Distribution of Bayes factors across the genome (B). Genomic landscape of scaffold 78 and 60 through bayes factor, F_ST_, shared polymorphism (ss) and diversity (π) (C). Data are from (Poelstra et al., 2014)

We next applied RIDGE on a genome-wide dataset produced for three pairs of *Corvus* species that form hybrid zones (pair RX: *C. corone* - *C. cornix*; pair XO: *C. cornix* - *C. orientalis*; pair OP: *C. orientalis* - *C. pectoralis*) where current gene flow is detected (Vijay et al., 2016).

The goodness-of-fit of the demographic parameters inferred by RIDGE was similar across all three pairs (RX: 0.33; XO: 0.21; OP: 0.26). The ratio of *T̂_split_* / 2 *N̂_e_* was approximately 0.5 for all three pairs (RX: 0.63; XO: 0.54; OP: 0.53) (Table S5), suggesting a comfort zone for RIDGE to detect gene flow barriers in all three datasets.

PCA analyses colored by BF show a first group of outliers (characterized by elevated levels of divergence, *F_ST_* and *Da*, and reduced level of diversity in all four pairs, Figures 7, 8 & S12). Those signals were consistent with theoretical expectations for gene flow barriers (i.e increased *Dxy*, *Da*, *sf*, *F_ST_*, and reduced ss and diversity). A second group of outliers, present in RX and XO pairs, displayed moderate increase in divergence but also in diversity and Tajima’s D, which corresponded to a more complex signature of gene flow barrier (Figure 8 & S12). In each pair, we identified a subset of loci with elevated Bayes factors (*BF* >50) clearly separated from the genome-wide distribution (Figure 8C). These subsets detected on a per locus basis (RX: 4.7%; XO: 0.37%; OP: 0.30%), represented smaller proportions than the expected proportion estimated in the general model *Q̂*(RX: 4.9%; XO: 4.8%; OP: 5.3%) but still fell within the credibility intervals (Figure 8B & Table S5).

**Figure 8:**
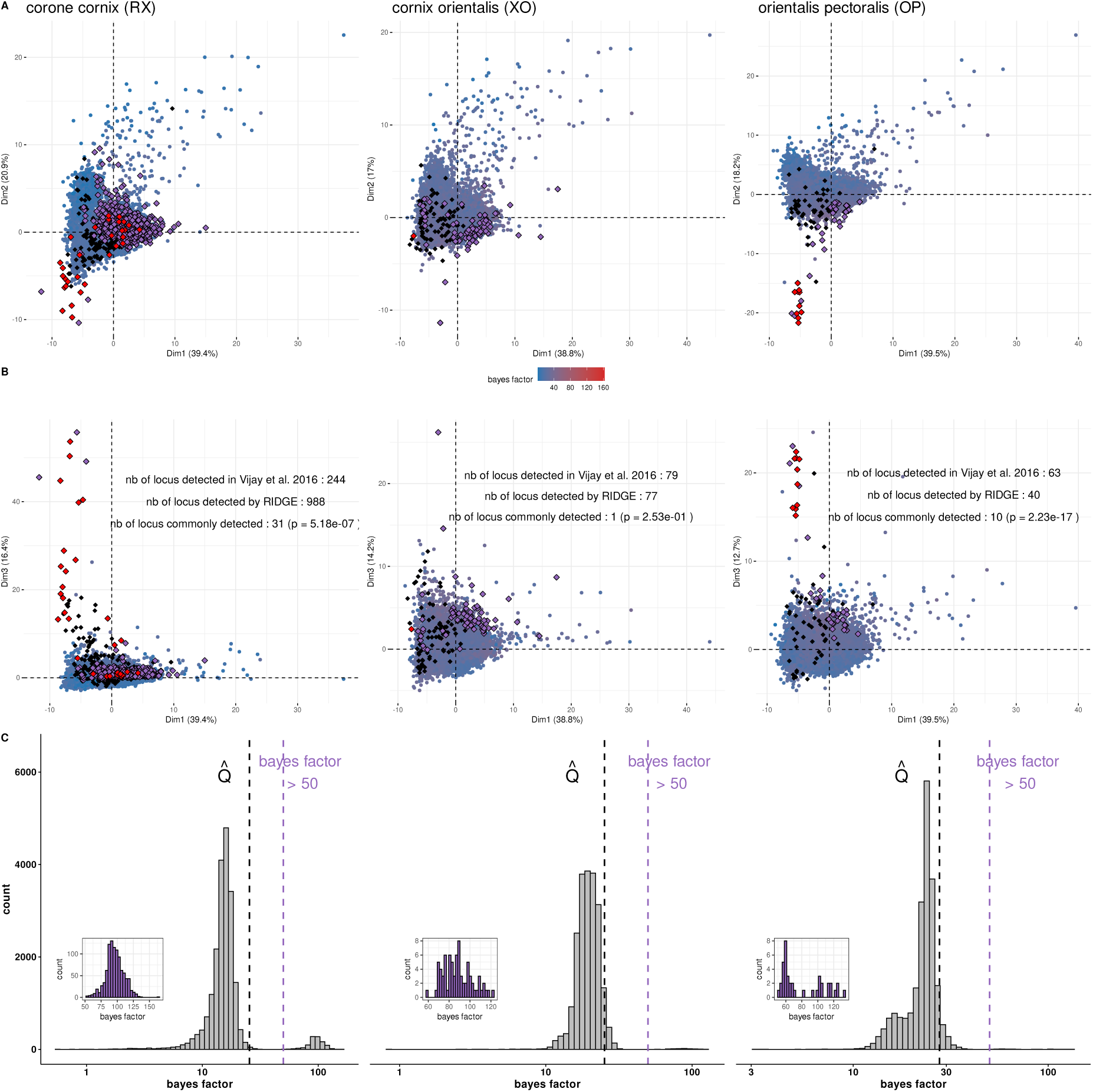
Barrier loci detection by RIDGE on three crow hybrid zones. PCA computed on summary statistics obtained from 50kb-windows along genomes with axes 1 and 2 (A) and 1 and 3 (B) displayed. Datapoints (windows) are colored according to the values of Bayes factors. Black diamonds represent loci detected in (Vijay et al., 2016), violet diamonds indicate loci detected by RIDGE that exceeded the population-specific Bayes factor threshold, and red diamonds represent loci detected both in (Vijay et al., 2016) and RIDGE. Distribution of Bayes factor values for each species pair (C). The histogram inside the figure shows the Bayes factor distribution of detected loci, which are the loci exceeding the population-specific Bayes factor threshold indicated by the violet dashed line. Black dashed line indicate the Bayes factor threshold based on the estimated barrier proportion Q̂. Data are from (Vijay et al., 2016).

We found significant overlap between our outliers and those of Vijay et al (2016) for the RX and OP pairs (Figure 8A & B). For OP, however, common outliers were found exclusively in the first group of outliers, whereas for RX common loci were found in the first and second group of outliers. On average, the BF revealed various correlation patterns among the three pairs, ranging from a clear correlation pattern with divergence statistics in the OP pair to a more blurred and complex pattern in the RX pair (Figure 9).

**Figure 9:**
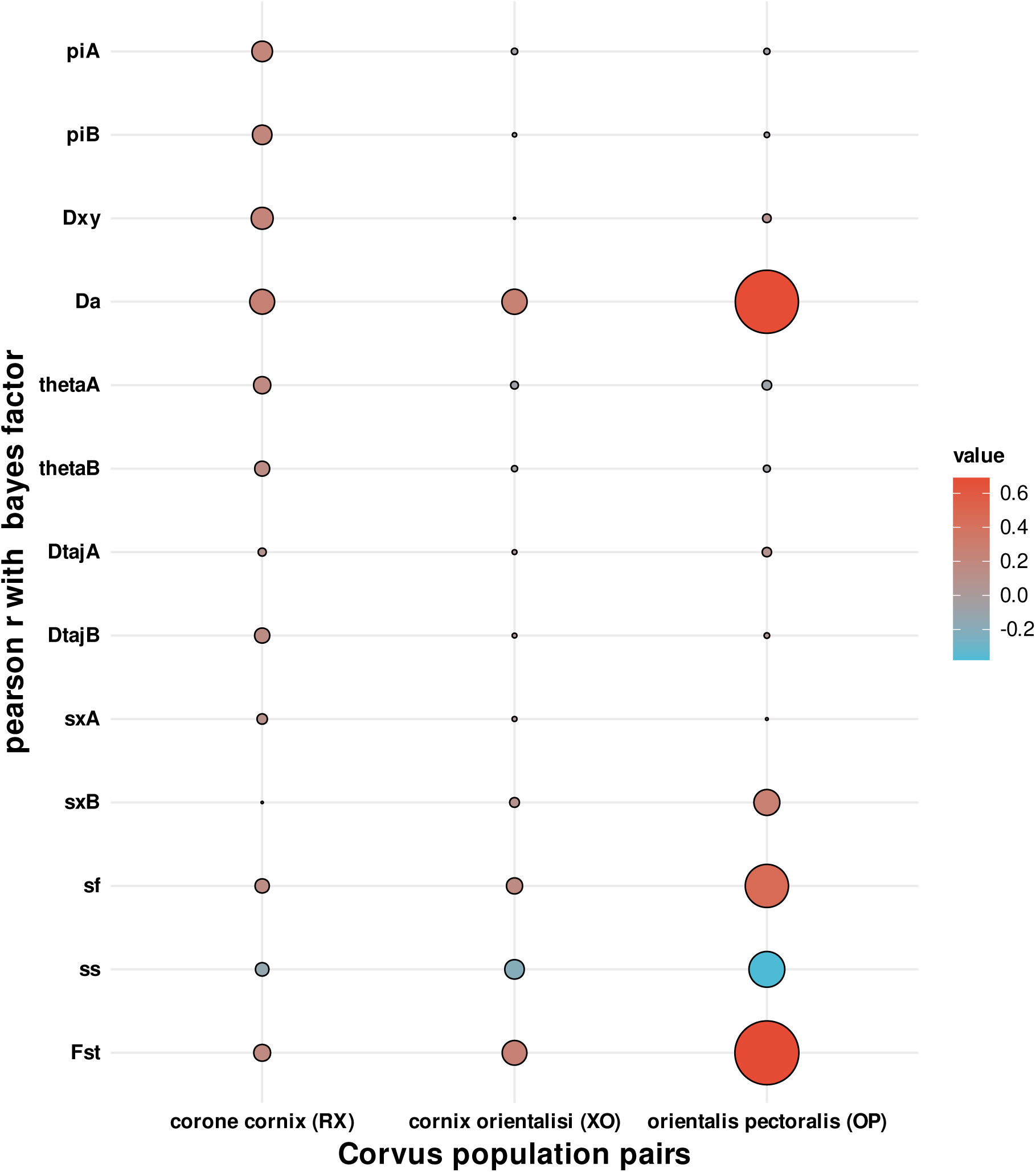
Pearson correlation between RIDGE Bayes factor and summary statistics used in the gene flow barrier detection for the three hybrid zones. Colors correspond to the values of correlations while circle size reflects the absolute values. Data are from (Vijay et al., 2016).

## Discussion

A key goal of speciation research is to elucidate the genetic mechanisms behind reproductive isolation. Although diverging populations have been analyzed in many studies, a challenging aspect remains the ability to capture the sequence of events that lead to the establishment of reproductive barriers. To answer this question, one approach is to compare populations that exhibit varying degrees of temporal and/or spatial divergence, including recently diverged ones. This requires the use of a comparative framework capable of detecting barriers to gene flow at both early and ancient stages across diverse biological systems, independently of their demographic history. In this context, we introduce RIDGE, a tool designed to facilitate this task.

### RIDGE offers a comparative framework where current migration is well captured

Currently, two methods explicitly model heterogeneity in the effective migration rate across the genome. Both tools utilize variations in effective population size to approximate selective effects along the genome. DILS (Fraïsse et al., 2021) uses an ABC framework under four demographic models of divergence (SI, IM, SC, AM) to assess alternative models of effective migration’s homogeneity/heterogeneity and provides corresponding genome-wide estimates. While not primarily designed to perform barrier detection, DILS can still provide valuable insights on potential barrier loci, conditioned on the selected demographic model (Fraïsse et al., 2021). There are however two main limits to this approach. Firstly, selecting a model can be rather arbitrary when two models explain the data equally well, which is often the case when divergence is shallow between populations (as shown in Fraïsse et al (2021) and confirmed here, Figure 3). Secondly, the use of potentially different demographic models complicates comparison across species pairs. gIMbl (Laetsch et al., 2023) relies on composite likelihood to identify windows of unexpected level of effective migration along the genome, but only under the IM model, while secondary contacts may be rather frequent in nature (ex: Leroy et al., 2020; Roux et al., 2013; Vijay et al., 2016)

RIDGE builds on DILS, offering a high degree of model flexibility, while proposing a comparative framework. In order to do so, RIDGE employs a model averaging approach by assigning weights to each demographic x genomic model without directing the user’s choice towards a single model. In addition, model averaging is also useful in reducing the uncertainty on parameter estimation when individual models present high variance (Dormann et al., 2018). Our results show that model averaging is especially relevant when data offers little discriminant power. For example, when *T_split_* is low, the discriminatory power of summary statistics is reduced, resulting in similar assignation to all models (Figure 3). Opting for the best scenario under such conditions might be misleading. For example, at *T_split_*=0.1∗2 *N_e_*, when current migration is simulated (IM or SC models), it is detected in only ~60% of the cases (Figure 3), thus potentially leading to the selection of the SI or AM models, thereby impeding the estimation of gene flow barriers. In contrast, the model averaging approach always provides an estimate of the proportion of gene flow barrier with a credibility interval, which can be large and include 0 when the statistical power is low. RIDGE thus allows for formal comparison of any datasets despite differences in demographic history and/or statistical power.

A direct consequence of using a demographic x genomic hypermodel is that RIDGE is not intended for precise estimation of a demographic model and its underlying parameters but rather to address demography as a confounding factor in the detection of gene flow barriers. High and stable values of goodness of fit across models and conditions indicate that we achieved this goal (Figure 2 & S1) and more moderately for complex/real scenario as for crows datasets (Table S5) where the goodness-of-fit is lower (*G_post_* ~ 0.9 for simulated datasets, *G_post_* ~ 0.25 for crows datasets). However, as expected, the accuracy of parameter estimation largely depends on the divergence time (Figure S4-S7). Similar to DILS (Fraïsse et al., 2021), the correct model’s contribution to parameter estimation and the detection of ongoing migration increases with divergence time (Figure 3). Overall, current migration is well captured, both in model weights and in parameter estimation (Figure 3, Figure S5).

This is well illustrated with the analysis of the crow datasets. After the ice cap had retreated in Europe around 10,000 years ago (~ 2000 crow generation), the ancestors of remnant carrion (*C. corone*) and hooded crow (*C. cornix*) populations met in a secondary contact in Central Europe, forming a narrow and stable hybrid zone (Knief et al., 2019; Metzler et al., 2021; Poelstra et al., 2014). Based on the sampling by Poelstra et al (2014), which covers a wide geographic area away from the central European hybrid zone, RIDGE favored the correct scenario, especially the occurrence of ongoing migration (model weight for SC = 48% and IM=41%) (Table S6). With the hybrid zone-specific dataset (RX pair), RIDGE encountered more difficulty in distinguishing between IM and SC scenarios, with IM at 49% and SC at 48%, likely due to the high levels of gene flow within the hybrid zone, which may have blurred the evidence of ancestral isolation to a greater extent than observed with the other sampling scheme (Poelstra et al., 2014). Overall, in all four datasets the current status of migration has been correctly captured with ongoing migration accounting for the majority of the model weight (RX: 94%; XO: 86%; OP: 86%; (Poelstra et al., 2014) : 90%).

### Informative summary statistics are highly context-dependent

One drawback of the ABC approach is that parameter inference relies on summary statistics to capture the genomic signal. Historically, *F_ST_*, a measure of relative divergence, has been the most widely used statistic in genome scans (Wolf & Ellegren, 2017). To avoid the confounding effect of reduced diversity in either of the compared populations due to other causes than barrier to migration (Cruickshank & Hahn, 2014; Ravinet et al., 2017), it is now common practice to combine it to absolute measure of divergence (*Dxy*) to other related statistics such as net divergence (*Da*) or the number of fixed differences (*sf*) (Han et al., 2017; Hejase et al., 2020). Here, we devised a new set of summary statistics based on outlier detection, and proved them to be useful for estimating barrier proportions. The reasoning was that loci showing local increase in divergence (measured by *F_ST_*, *Dxy*, *Da*, *sf*) and decrease in diversity would generate outliers in the genome wide divergence and diversity distributions. Our results show that outlier statistics mostly contribute to *Q̂*under moderate gene flow (*M* =1), and mainly for low level of barrier proportion (*Q* <0.1) (Figure S9) where estimation of barrier proportion may be challenging.

An important result of our study is that the set of summary statistics that effectively capture the signal of barrier loci varies with the divergence and demographic history and with the sampling scheme (Figure 9). In the OP and XO pairs, Bayes factor outliers are mainly captured by *F_ST_*, *ss* and *Da* statistics (exposing *F_ST_* and *Da* increase, *ss* reduction and also moderate reduction of diversity) with a stronger signal in OP than XO (Figure S12). For the remaining outliers in PO and XO and for all outliers in the RX pair, in addition to an expected increase in divergence, outliers show a moderate increase in diversity statistics, which is the genomic pattern theoretically expected for a gene flow barrier evolving under low-intensity divergent selection which generates an excess of maladaptive alleles and thus increases diversity (Sakamoto & Innan, 2019). Differences between correlation patterns between summary statistics and BF could reflect the difference in the environment in which incipient crows species evolved, but also the difference in the geographical area covered by the hybrid zone (Vijay et al., 2016).

These examples illustrate that considering a few statistics in the detection of barrier loci can be misleading as signatures can be complex and context-dependent. It thus advocates for the use of a more inclusive approach as implemented in the BF derived from the random-forest-based ABC approach of RIDGE. One contribution of the Random Forest (RF) is to reduce the curse of dimensionality (Bellman & Kalaba, 1959), which improves accuracy and computation time, RF also makes ABC a calibration-free problem by automating the inclusion of summary statistics (Raynal et al., 2019). In return, a possible drawback is that RF results are less interpretable due to their complex nature. Indeed, even if the *abcrf* package provides a way to understand the contribution of variables to parameters estimations, it still remains difficult to interpret the RF decision for a specific locus.

### Detection of barrier loci using RIDGE

We validated the ability of RIDGE to detect gene flow barriers on empirical datasets from Poelstra et al (2014) and Vijay et al (2016). In particular, we clearly detected the large and well-established region of scaffold 78 on chromosome 18. It contains major loci that are involved in mate choice patterns between *C.corone* and *C.cornix* (RX) (Knief et al., 2019; Metzler et al., 2021; Poelstra et al., 2014). The study by (Vijay et al., 2016) was conducted on three species pairs that had similar demographic histories. For all three pairs of populations, we identified a portion of loci exhibiting elevated BF. We found significant overlap between our results and previously detected outliers for the RX and OP pairs. The overlap mainly corresponded to extreme outliers – characterized by highly divergent loci between species and reduced diversity. We also identified loci not previously detected (Vijay et al., 2016). Those likely corresponded to barriers evolving under low intensity divergent selection as they displayed both increased divergence and diversity (Sakamoto & Innan, 2019). Conversely, barrier loci detected in Vijay et al (2016) but not with RIDGE display low diversity without distinctive divergence patterns. This observation can be attributed to the confounding effect of the heterogeneity in *N_e_*, not explicitly accounted for in Vijay et al (2016) and which is a classic pitfall of *F_ST_*scan approaches (Cruickshank & Hahn, 2014). The fact that RIDGE detected only a limited number of loci displaying such a pattern implies that it effectively circumvents this problem. For XO pair, due to its spatial range (three to seven times wider than the hybrid zone of RX pair), selection strength is reduced (Vijay et al., 2016), resulting in candidate regions showing low-intensity divergence patterns in Vijay et al (2016) results similarly to our results (Figure 8). Furthermore, since low signal can increase noise in detection results, we did not detect any direct overlap between the candidate XO gene from Vijay et al. (2016) and our results. However, when examining the regions surrounding the candidate gene, we observed common regions such as the gene *LRP5*, which was consistently present in XO and OP pairs in Vijay and was consistently located at a distance of 50-100 kb from an outlier locus in our results.

### Benefits of RIDGE and Guidelines for its use

RIDGE relies on an ABC approach that offers a lot of flexibility, enabling it to explore genomic heterogeneity and to incorporate customized summary statistics. We have also devised a method for generating multidimensional parameter estimates, extending beyond the initial single-parameter focus of *abcrf* (Raynal et al., 2019). This improvement enables RIDGE to deal effectively with parameter interdependencies and increase the precision of parameter estimations. Another improvement introduced by RIDGE is the incorporation of Bayes factors, facilitating result comparisons.

The simulated datasets we explored gave us guidelines for the conditions where RIDGE can provide useful and accurate results. We suggest to use datasets with SNP density higher than 0.1%, such as in crows and simulated datasets, where the SNP density was around 1%. We also advise to use a minimum of three samples per population. The goodness-of-fit statistics enables users to check the quality of inferences made. If *G_post_* < 5%, the user should verify the prior bounds. The guidelines for interpreting and thresholding BF depend on the user’s goals. If RIDGE is used solely to discover new candidate genes involved in gene flow barriers for a specific population pair, we recommend using a customized threshold that optimally captures Bayes factor outliers. For the purpose of comparison, it is recommended to use a standard threshold for all datasets, for example *BF* >100 or to keep the number of outlier loci corresponding to the proportion of barriers estimated in the first step of RIDGE (*Q̂*). Crucially, genomic data alone cannot provide conclusive evidence of barrier loci and so RIDGE results should be coupled with other analysis such as functional analysis (Ravinet et al., 2017). It is worth noting that window length (default set to 10 kb) can significantly affect the results of RIDGE. It should be determined according to the extent of linkage disequilibrium as well as the level of diversity, since it determines the amount of polymorphism and consequently affects the strength of the signal.

As is the case with all ABC approaches, the quality of the priors given by the user affects the results obtained using RIDGE. A *T_split_* of 0.1*2*N_e_* generations (10,000 generations in our simulations) appears to be a lower bound for both demography (Fig. 4 & 5) and barrier inferences (Fig. 6), below which RIDGE fails to capture informative signals. RIDGE can detect gene flow barriers on both simulated (Fig. 6) and empirical data (Fig. 7), starting at 0.1*2*N_e_* generation, which represents a very low level of divergence. For context, DILS correctly inferred a gene flow barrier when *T_split_* >0.5 2*N_e_* generations, while gIMbl demonstrated its effectiveness on one pair of *Heliconius* species that diverged 4.5 million generations ago, estimated to represent 0.49 x 2*N_e_* generations (Martin et al., 2015).

Comparative approaches have been useful in understanding the genomic basis involved in the process of reproductive isolation (e.g (Roux et al., 2016)) and they will continue to play an important role in speciation research. By its flexibility and its comparative framework, RIDGE should become a useful tool to follow this direction.

## Acknowledgement

We thank Camille Roux for the help with the DILS code and Miguel de Navascués for advice in the use of the ABC-RF method. We also thank Thibault Leroy, Christelle Fraïsse, Yves Vigouroux, Maxime Bonhomme and Claire Mérot for their insightful discussions and valuable inputs during the course of the project. We thank Augustin Desprez, Harry Belcram, Clemetine Tocco and Arthur Wojcik for helping to improve RIDGE by beta-testing it. We would also thank Chyi Yin Gwee and Jochen Wolf for providing us with the pre-mapped VCF dataset of crows. This work benefited from the computing resources provided by the GenOuest cluster,the Cornuta cluster, and the IFB core cluster.

This work was supported by the grant DomIsol overseen by the French National Research Agency (ANR-19-CE32-0009-02). GQE-Le Moulon benefits from the support of Saclay Plant Sciences-SPS (ANR-17-EUR-0007) as well as from the Institut Diversité, Ecologie et Evolution du Vivant (IDEEV). E.B. was financed by a doctoral contract from DomIsol and from Région Bretagne through the Doctoral School EGAAL. In addition, E.B. benefited from a travel grant from GDR 3765 “Approche Interdisciplinaire de l’Évolution Moléculaire”.

## Data accessibility statement

Source codes to deploy RIDGE and user manual are freely available from GitHub: https://github.com/EwenBurban/RIDGE.git. This GitHub repository also includes a pipeline for simulating pseudo-observed datasets and an optimized pipeline for running RIDGE on thousands of pseudo-observed datasets.

## Author Contributions

designed research: Maud Tenaillon, Sylvain Glémin

performed research: Ewen Burban

Funding acquisition: Maud Tenaillon, Sylvain Glémin

informatics tool development: Ewen Burban

analyzed data: Ewen Burban

supervision: Sylvain Glémin, Maud Tenaillon

wrote the paper - original draft : Ewen Burban

wrote the paper – review & editing: Ewen Burban, Sylvain Glémin, Maud Tenaillon

