## Supplementary_material for "RIDGE, a tool tailored to detect gene flow barriers across species pairs"

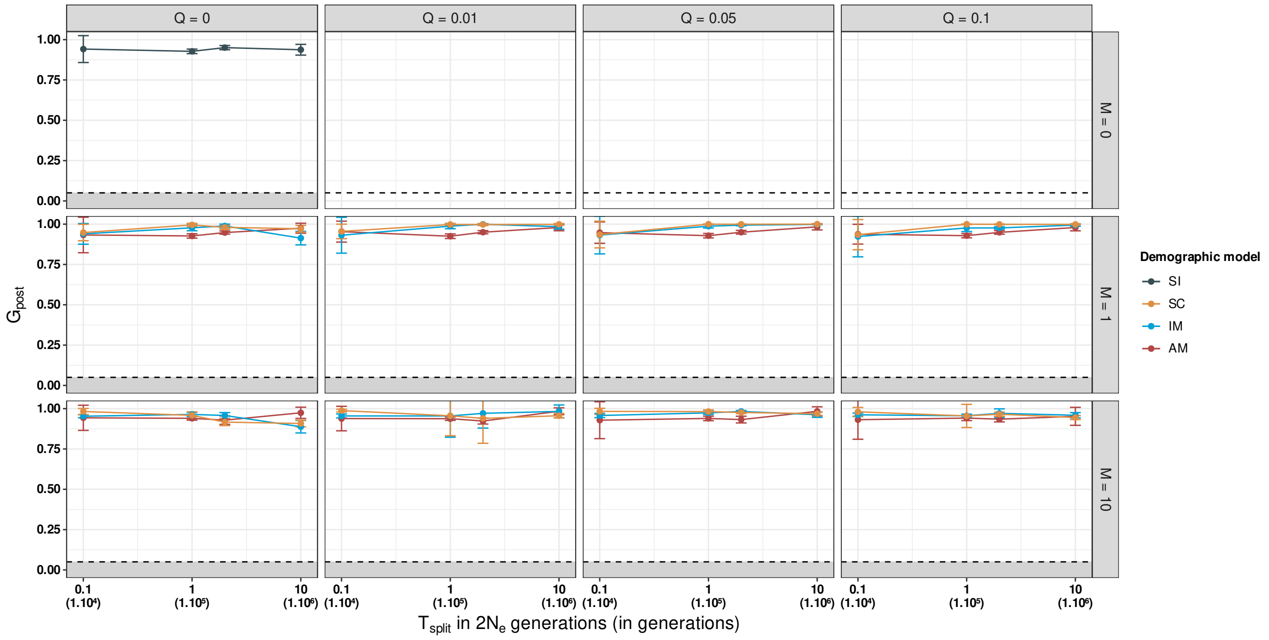


*Figure S1: Evolution of the goodness-of-fit of the posteriors as a function of $T_{split}$, migration ($M$) and barrier proportion ($Q$), for four demographic models. The gray zone represents the rejection zone, in which inferred models are discarded. Average values over 100 replicates with error bars (standard deviation) are presented. Pseudo-observed datasets were simulated under 2M2N and 1M2N models.*

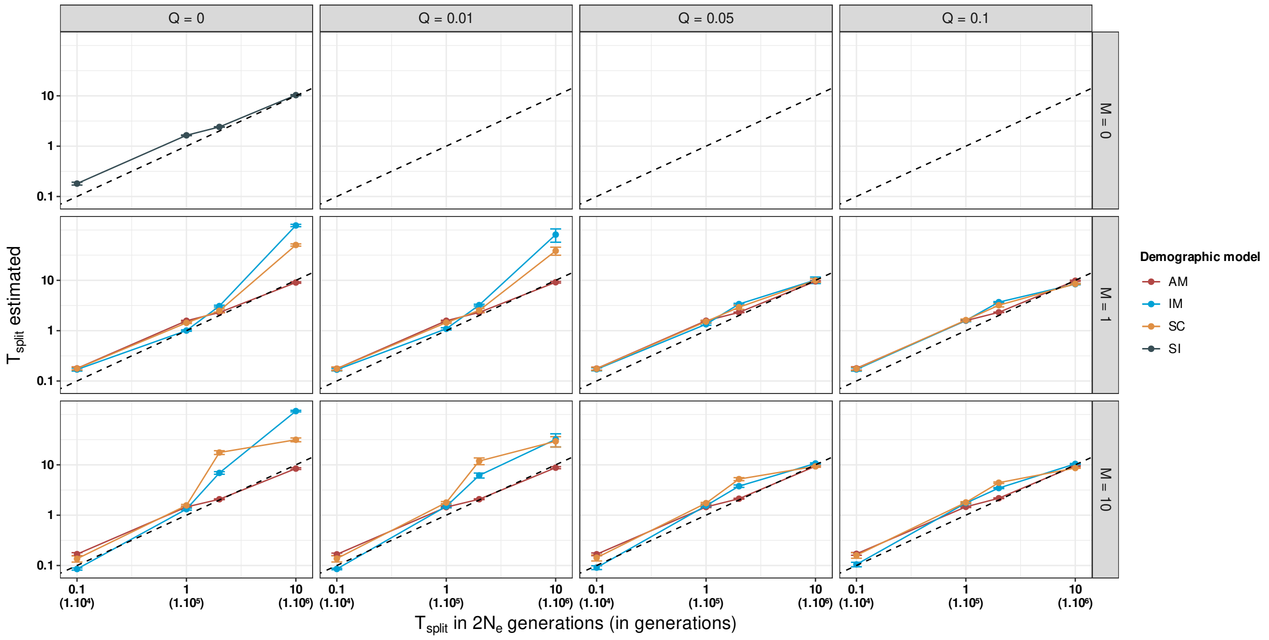


*Figure S2: Estimated Time split ($\hat{T}_{split}$) as a function of simulated time ($T_{split}$) split, migration ($M$) and barrier proportion ($Q$) for four demographic models. Average values over 100 replicates with error bars (standard deviation) are presented. Pseudo-observed data were simulated under 2M2N and 1M2N. The dashed line represents the reference (simulated = estimated).*

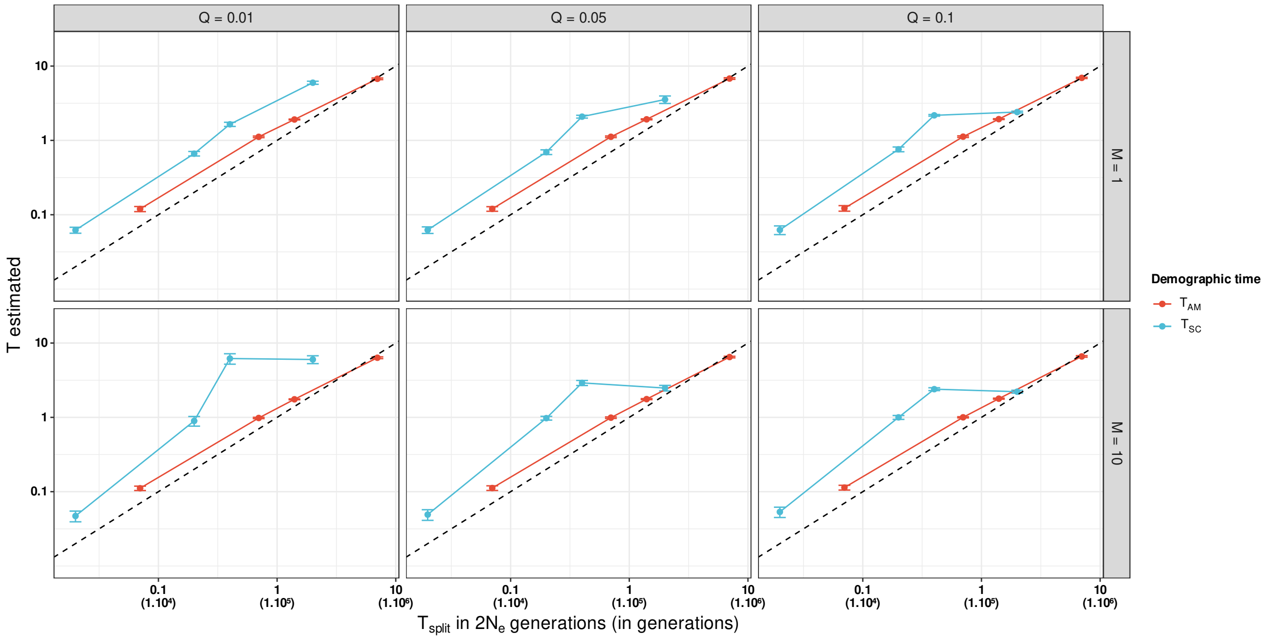


*Figure S3: Estimated Time of secondary contact ( $\hat{T}_{SC}$ ) and time of last migratory contact ( $\hat{T}_{AM}$ ) as a function of simulated time split ( $T_{split}$ ), migration ( $M$ ) and barrier proportion ( $Q$ ) for respectively SC and AM demographic model. Average values over 100 replicates with error bars (standard deviation) are presented. Pseudo-observed data were simulated under 2M2N. The dashed line represents the reference (simulated=estimated).*

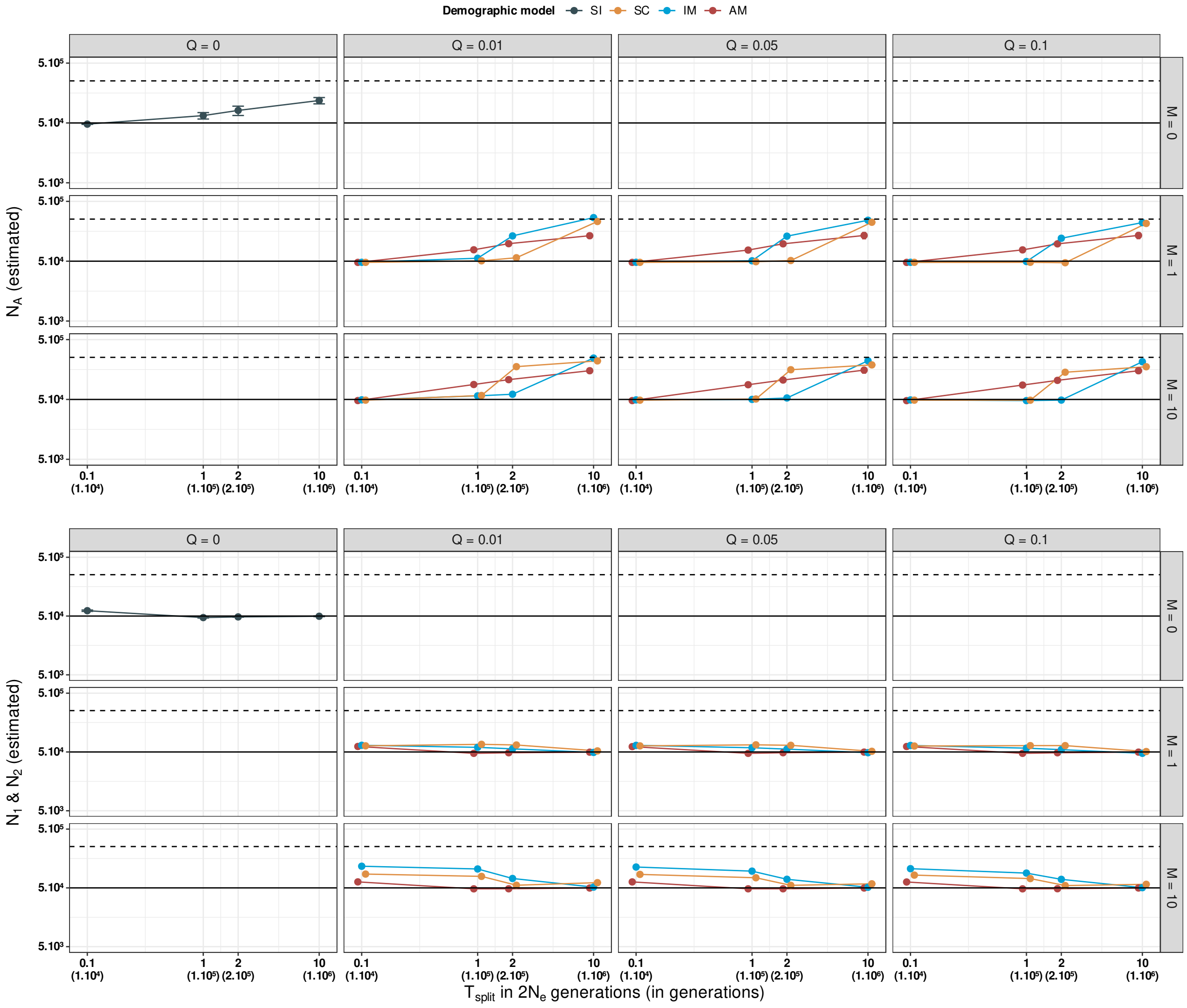


*Figure S4: Estimated population size (past with $\hat{N}_A$ and current with $\hat{N}_1$ and $\hat{N}_2$) under fours demographics models. Average values over 100 replicates with error bars (standard deviation) are presented. The plain line represents the value used in the simulation ($N_e$ = 50 000), and the dashed line represents the mean value of priors ($N_e$ = 252 500). Simulated data were obtained under 2M2N.*

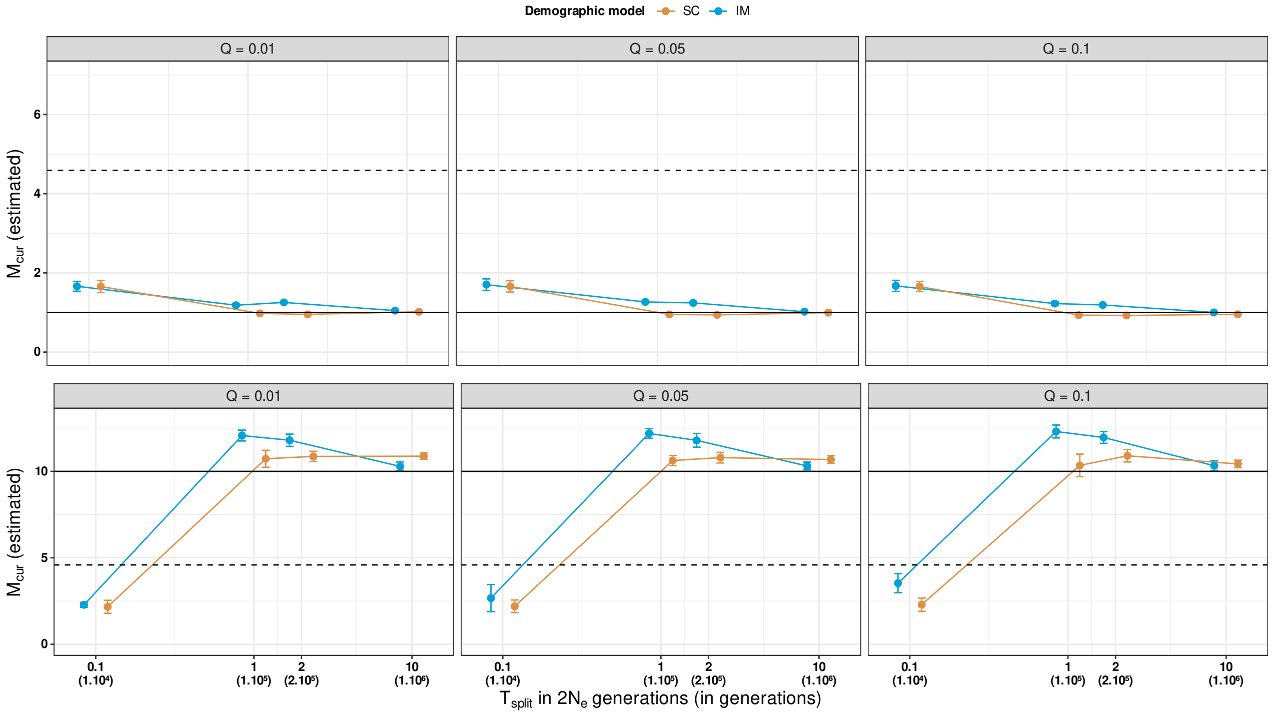


*Figure S5: Current migration rate ( $\hat{M}_{cur}$ ) estimation accuracy under IM and SC models under 2M2N.* Average values over 100 replicates with error bars (standard deviation) are presented. *The plain black line represents the true value ( $M_{cur}$ =1 and 10) used to generate the pseudo-observed datasets and the dashed line represents the mean of priors* $M = 4.58$ .

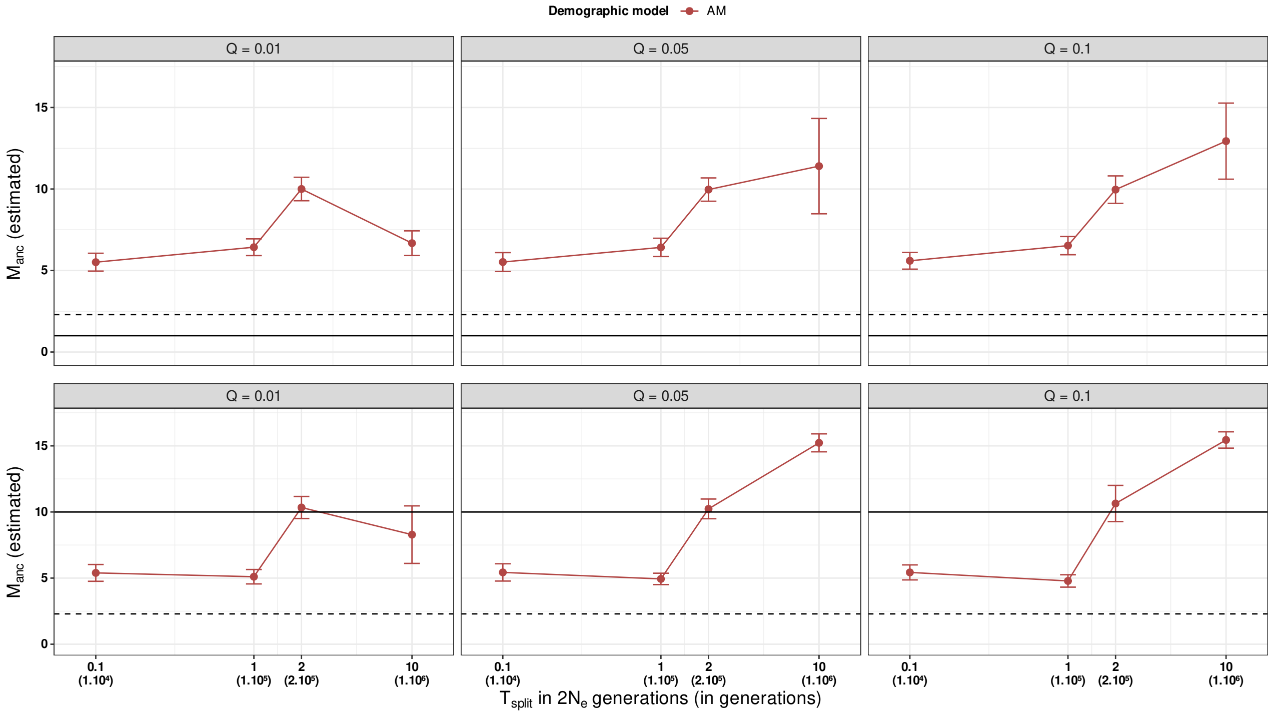


*Figure S6: Ancestral migration rate ( $\hat{M}_{anc}$ ) estimation accuracy for AM model under 2M2N. Average values over 100 replicates with error bars (standard deviation) are presented. The plain black line represents the true value ( $M_{anc}$ = 1 and 10) used to generate the pseudo-observed datasets and the dashed line represents the mean of priors* $M = 2.29$ .

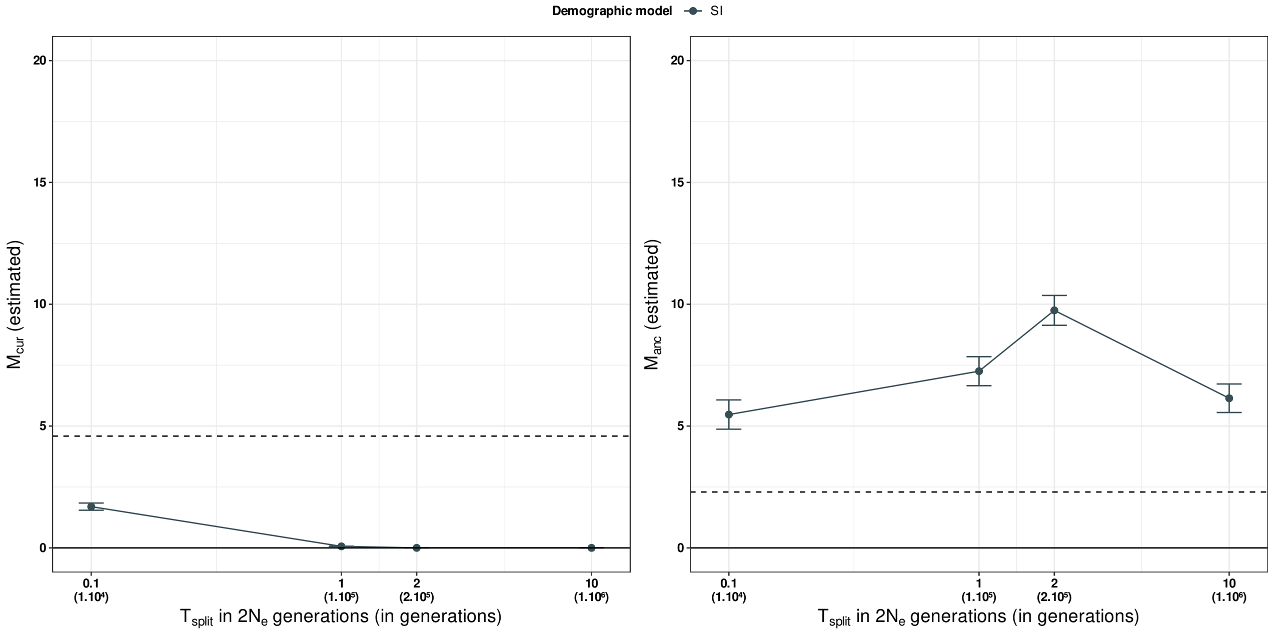


*Figure S7: Current (left) and Ancestral (right) migration rate estimation accuracy under SI model. Average values over 100 replicates with error bars (standard deviation) are presented. The plain black line represents the true value ( $M_{cur} = M_{anc} = 0$ ) used to generate the pseudo-observed datasets and the dashed line represents the mean of priors $M = 4.58$ (left) and $M = 2.29$ (right).*

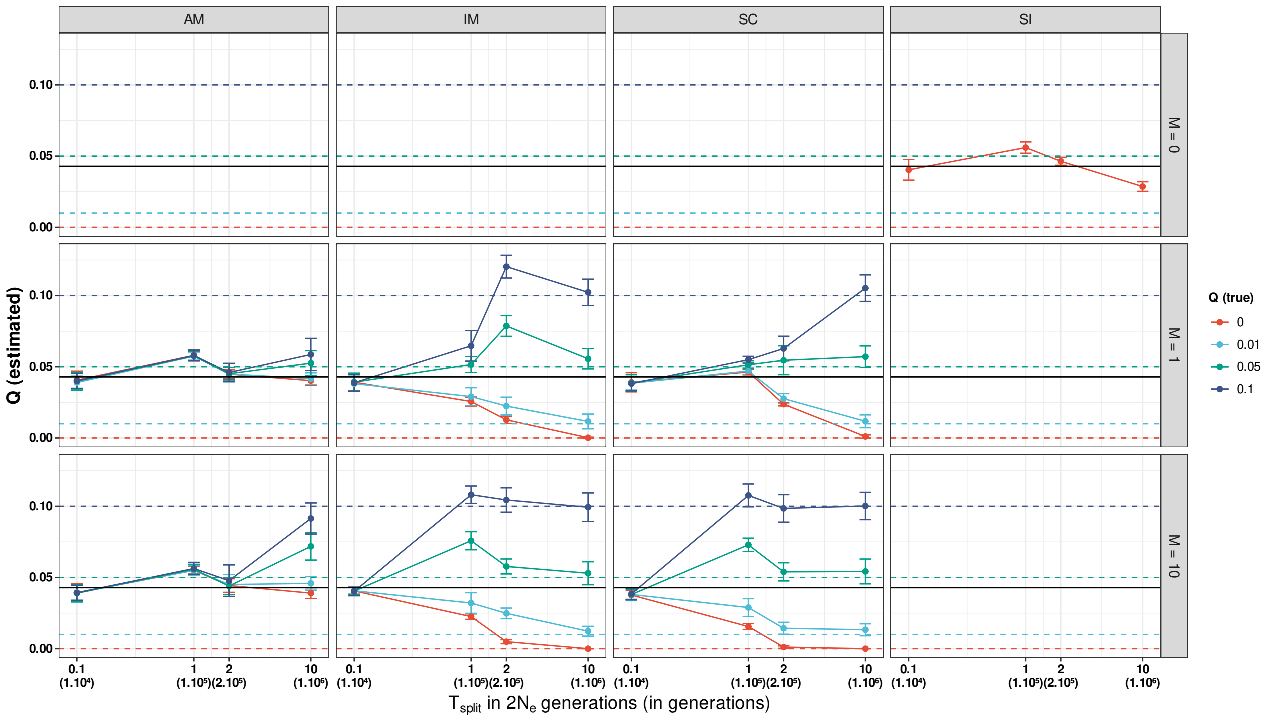


*Figure S8: Barrier proportion estimates ( $\hat{Q}$ ) as a function of divergence time under four demographic models. Average values over 100 replicates with error bars (standard deviation) are presented and the plain black line represents the mean of priors $Q = 4.2\%$ . Dashed lines represent references values corresponding to each barrier proportion conditions.*

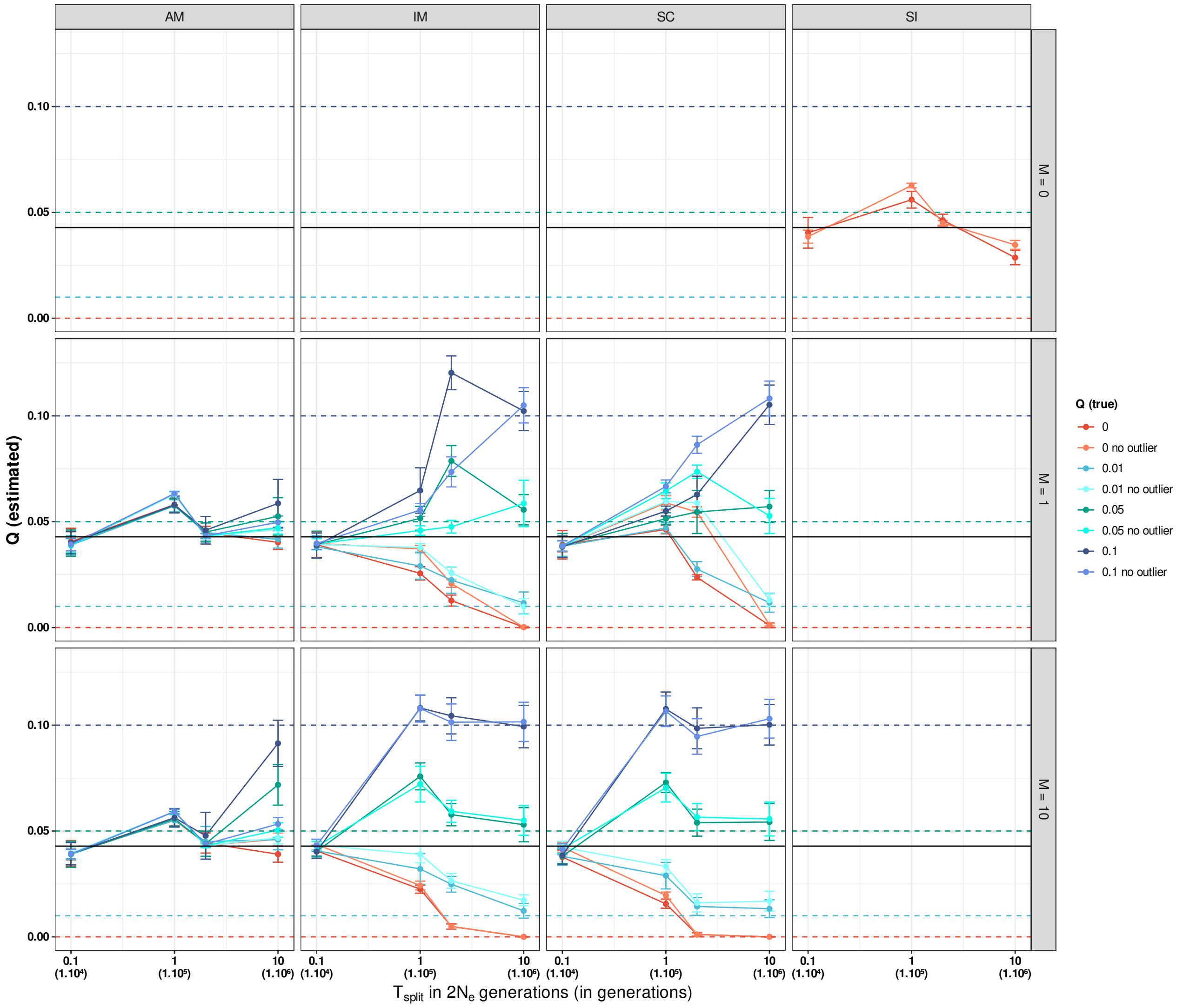


*Figure S9: Comparison between barrier proportion estimates with or without outlier summary statistics (no outlier) as a function of divergence time under the fours demographic models. Average values over 100 replicates with error bars (standard deviation) are presented and the plain black line represents the mean of priors* $Q = 4.2\%$ *. Dashed lines represent the initial value of barrier proportion used in pseudo-observed dataset.*

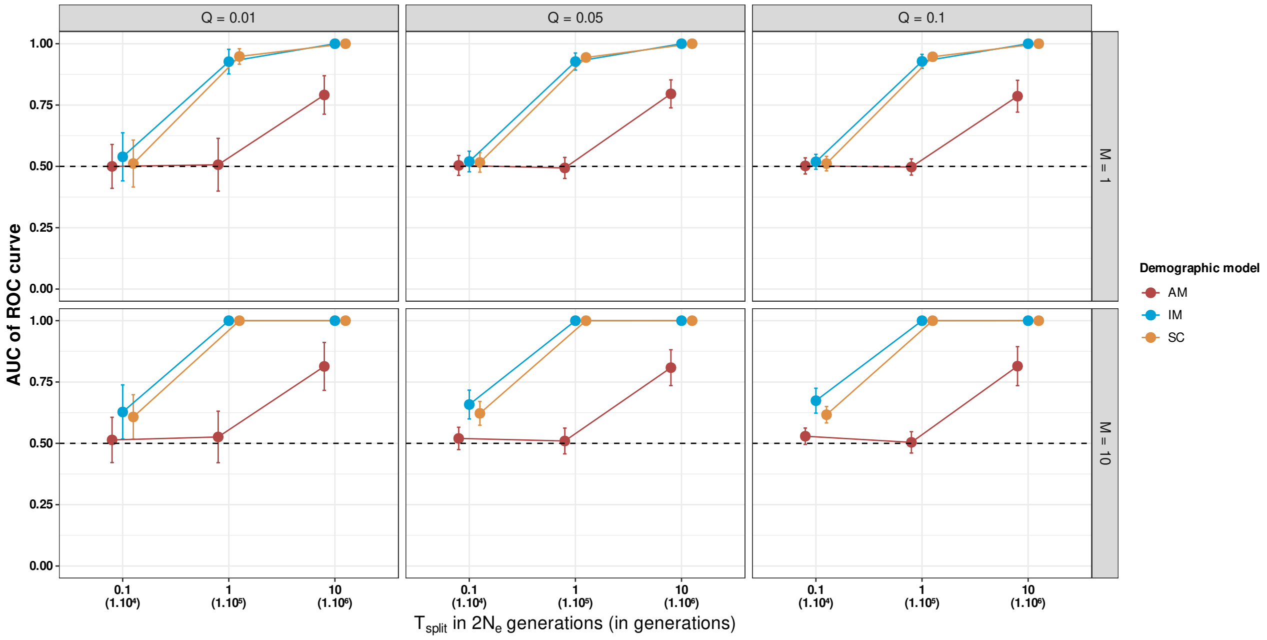


*Figure S10: Discriminant power measured through the AUC of ROC as a function of divergence time $T_{split}$, migration $M$, demographic model and the proportion of barrier $Q$. The AUC relates the False Positive Rate (FPR) to the True Positive Rate (TPR), the greater the AUC the higher the discriminant power . Average values over 100 replicates with error bars (standard deviation) are presented. The dashed line represents the AUC=0.5 threshold, above which signal is captured.*

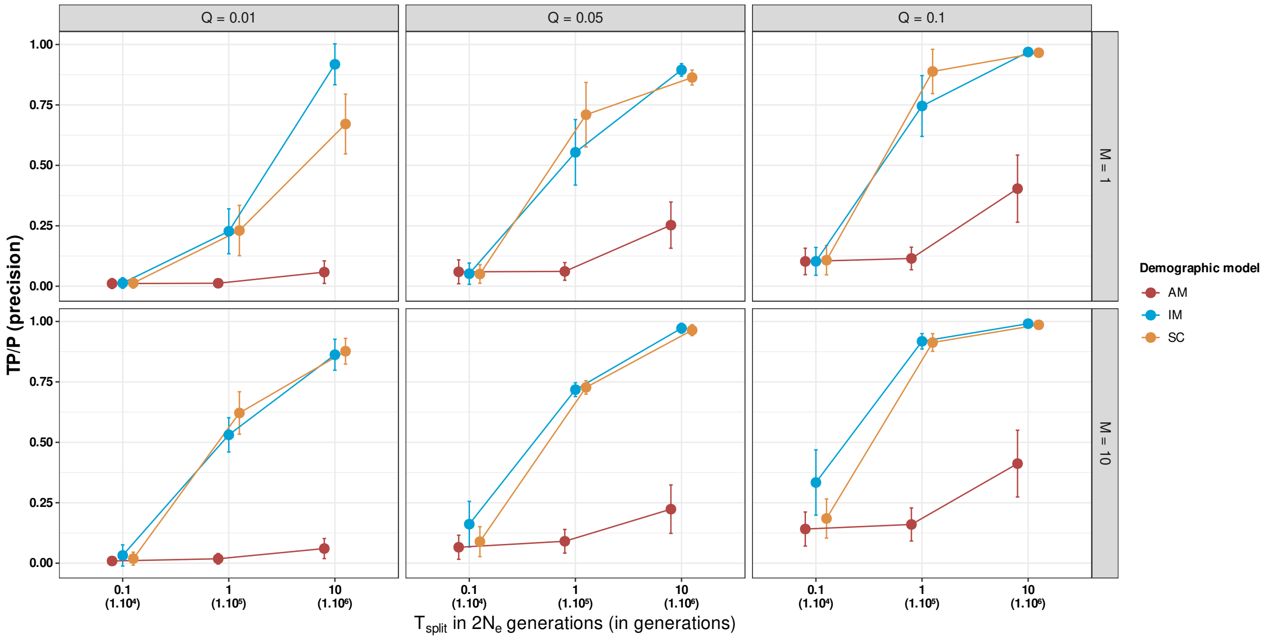


*Figure S11: Precision of barrier loci identification as a function of divergence time $T_{split}$, migration $M$, model and the proportion of barrier $Q$. Precision, which is the ratio of the number of true positives (TP) divided by the number of detected loci (P) – which is true positives plus the number of false positives – are shown with average values over 100 replicates with error bars (standard deviation) are presented. Detected loci are loci that exhibit a Bayes factor exceeding the threshold set to the quantile at $1-\hat{Q}$.*

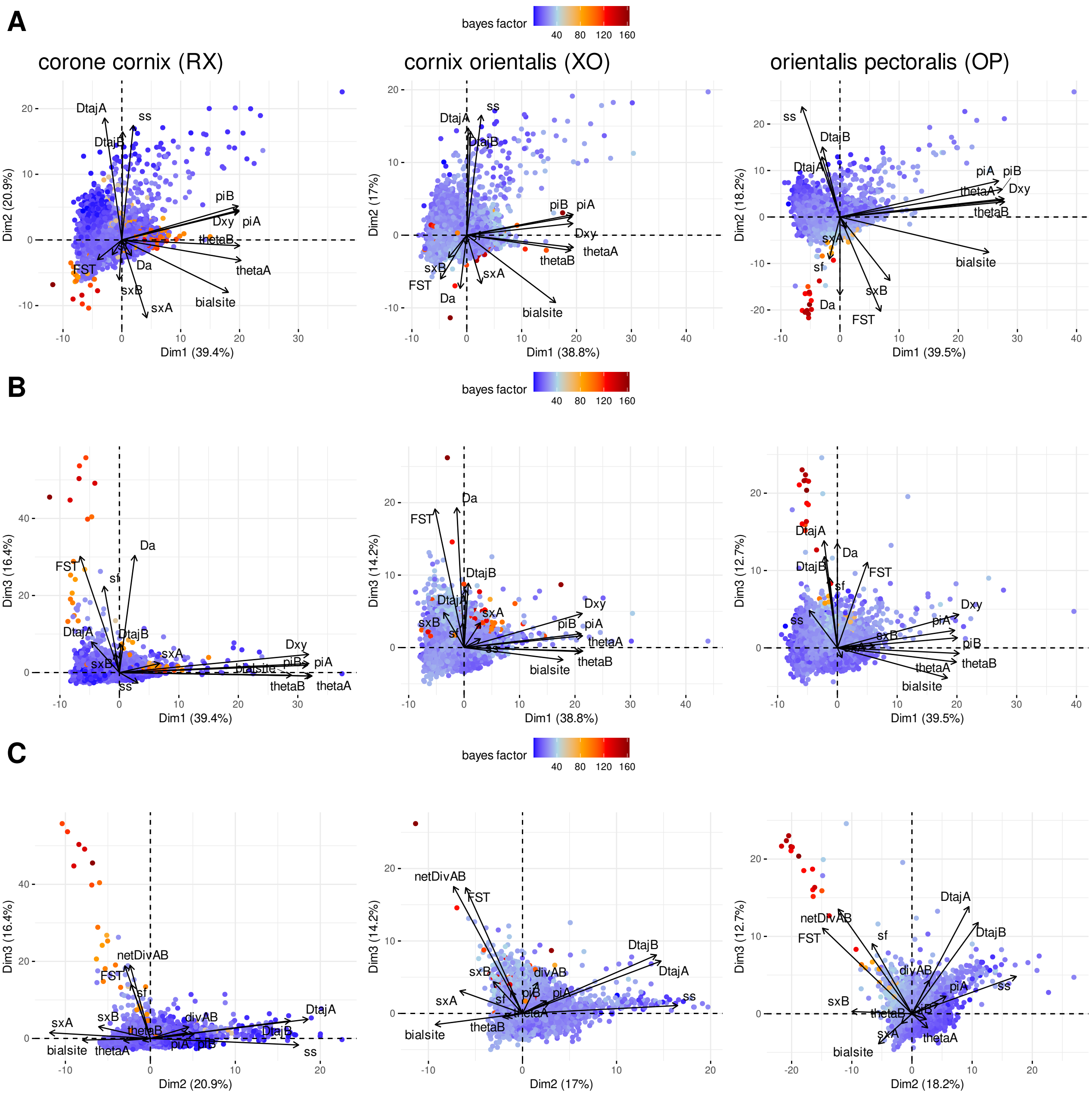


*Figure S12: PCA computed on summary statistics obtained from 50kb-windows along genomes with axes 1 and 2 (A) and 1 and 3 (B) and 2 and 3 (C) displayed. Data points (windows) are colored according to the values of Bayes factors. Data are from (Vijay et al, 2016).*

*Table S1: Demographic parameters used under four demographic models (SI: Strict Isolation, IM: Isolation Migration, SC: Secondary Contact, AM: Ancestral Migration) and four Genomic model (1M, 2M, 1N, 2N). Parameters are either estimated (empty field) or fixed to a value defined as indicated - either 0 or the value of another parameter. Note that for a single simulation, K value is drawn in U[0,1] only once, so it means that* $T_{SC}=T_{AM}=K*T_{split}$ .

| | AM | | | | IM | | | | SC | | | | SI | |
|---|---|---|---|---|---|---|---|---|---|---|---|---|---|---|
| | 1M1N | 1M2N | 2M1N | 2M2N | 1M1N | 1M2N | 2M1N | 2M2N | 1M1N | 1M2N | 2M1N | 2M2N | 1N | 2N |
| $T_{split}$ | | | | | | | | | | | | | | |
| $T_{AM}$ | | | | | $K*\ T_{split}$ | $K*\ T_{split}$ | $K*\ T_{split}$ | $K*\ T_{split}$ | $T_{split}$ | $T_{split}$ | $T_{split}$ | $T_{split}$ | $T_{split}$ | $T_{split}$ |
| $T_{SC}$ | 0 | 0 | 0 | 0 | $K*\ T_{split}$ | $K*\ T_{split}$ | $K*\ T_{split}$ | $K*\ T_{split}$ | | | | | 0 | 0 |
| *Na* | | | | | | | | | | | | | | |
| *N1* | | | | | | | | | | | | | | |
| *N2* | | | | | | | | | | | | | | |
| $M_{cur}$ | 0 | 0 | 0 | 0 | | | | | | | | | 0 | 0 |
| $M_{anc}$ | | | | | $M_{cur}$ | $M_{cur}$ | $M_{cur}$ | $M_{cur}$ | 0 | 0 | 0 | 0 | 0 | 0 |
| $\alpha$ | $1.10^4$ | | $1.10^4$ | | $1.10^4$ | | $1.10^4$ | | 1e4 | | $1.10^4$ | | $1.10^4$ | $1.10^4$ |
| $\beta$ | $1.10^4$ | | $1.10^4$ | | $1.10^4$ | | $1.10^4$ | | $1.10^4$ | | $1.10^4$ | | $1.10^4$ | $1.10^4$ |
| $Q_{anc}$ | 0 | 0 | | | 0 | 0 | $Q_{cur}$ | $Q_{cur}$ | 0 | 0 | 0 | 0 | 0 | 0 |
| $Q_{cur}$ | 0 | 0 | 0 | 0 | 0 | 0 | | | 0 | 0 | | | 0 | 0 |

*Table S2: Parameter values used in the simulations of pseudo-observed datasets. Note that for the strict isolation model, only* $T_{split}$ *varies. The number of loci in a pseudo-observed dataset is 1000 loci of 10 kb each. The mutation rate was set to* $1.10^{-8}$ *and the recombination rate to* $1.10^{-7}$ *event/generation/bp. Populations size* $N_1 = N_2 = N_A = 5.10^4$ *individuals. From each daughter's populations, 20 haploid samples are produced. Each condition is repeated 100 times. To run RIDGE on each pseudo-observed dataset, prior were defined as follows:* $T_{split}$ *and Ne prior distribution is bounded by one order below and above the true value (e.g, for* $1.10^5$ *the distribution is bounded between* $1.10^4$ *and* $1.10^6$*). The* $M$ *prior distribution is bounded between 0.1 and 50 N.m, and the* $Q$ *prior distribution is bounded between 0 and 0.2.*

| Parameter | Parameter values |
|---|---|
| *Tsplit* | $1.10^4$, $1.10^5$, $2.10^5$, $1.10^6$ |
| $M_{cur}$ and $M_{anc}$ | 1,10 |
| $Q_{cur}$ and $Q_{anc}$ | 0.01, 0.05, 0.1 |

*Table S3: Prior bound used to run RIDGE over all crow population pairs*

| Parameter | Parameter bound (min - max) |
|---|---|
| Tsplit | 10 000 - 150 000 |
| Ne (population size) | 30 000 - 250 000 |
| m | 0.1 - 50 |
| Q | 0 - 0.2 |

*Table S4: Pearson correlation (r) between estimated proportion of barrier $Q$ and outlier statistics under three demographic models with different Time of split. Simulations were ran under 2M2N model with $M=10$ and $Q=0$. Values of r >0.5 are shown in bold, NA indicates that correlation could not be computed.*

| Model | Tsplit | r between Q and outlier statistic | | | | |
|---|---|---|---|---|---|---|
| | | $F_{ST}$ | *Dxy* | *Da* | *sf* | $\pi$ |
| AM | $1.10^4$ | 0.27 | -0.15 | **0.68** | 0.19 | -0.15 |
| | $1.10^5$ | -0.49 | -0.27 | **0.56** | -0.37 | -0.15 |
| | $2.10^5$ | 0.16 | 0.15 | **0.74** | 0.04 | 0.09 |
| | $1.10^6$ | -0.21 | -0.09 | **0.70** | -0.15 | -0.21 |
| IM | $1.10^4$ | NA | -0.33 | NA | **0.77** | -0.07 |
| | $1.10^5$ | NA | 0.08 | NA | **0.97** | 0.04 |
| | $2.10^5$ | 0.12 | -0.21 | 0.12 | **0.99** | -0.21 |
| | $1.10^6$ | -0.08 | **1.00** | -0.08 | **1.00** | -0.20 |
| SC | $1.10^4$ | -0.08 | **0.68** | 0.36 | 0.08 | -0.08 |
| | $1.10^5$ | -0.02 | 0.03 | **0.97** | -0.15 | -0.02 |
| | $2.10^5$ | -0.07 | 0.03 | **0.99** | -0.06 | -0.07 |
| | $1.10^6$ | **1.00** | NA | **0.99** | -0.24 | **1.00** |

*Table S5: Estimated demographic and genomic parameters for each pair of crow species from Vijay et al (2016) and Poelstra et al (2014) (=Poelstra comp). For each parameter, the mean is presented with a credibility interval [5%;95%]. Note that time are expressed in crow generations, migration in N.m units and population size in number of individuals.*

| | RX | XO | OP | Poelstra comp |
|---|---|---|---|---|
| $\hat{N}_1$ | 159708<br>[42144;243218] | 150178<br>[39250;243379] | 157555<br>[47254;244924] | 151414<br>[45207;242718] |
| $\hat{N}_2$ | 143563<br>[39238;239724] | 157750<br>[43267;243150] | 149550<br>[42403;240966] | 155742<br>[49362;243728] |
| $\hat{N}_A$ | 107353<br>[37905;213268] | 110599<br>[36190;226339] | 109967<br>[34430;227025] | 96606<br>[42713;193155] |
| $\hat{M}_{cur}$ | 20.94<br>[0;46.08] | 16.13<br>[0;44.85] | 18.29<br>[0;46.18] | 13.69<br>[0;43.91] |
| $\hat{M}_{anc}$ | 10.66<br>[0;40.12] | 8.6<br>[0;40.42] | 10.66<br>[0;41.28] | 6.43<br>[0;36.31] |
| $\hat{\alpha}$ | 2093.07<br>[0.4;10000] | 1803.27<br>[0.31;10000] | 1913.1<br>[0.42;10000] | 2124.28<br>[1.16;10000] |
| $\hat{\beta}$ | 2094.17<br>[0.72;10000] | 1803.9<br>[0.63;10000] | 1913.83<br>[0.62;10000] | 2124.28<br>[1.03;10000] |
| $\hat{T}_{SC}$ | 27404<br>[0;85824] | 26761<br>[0;97143] | 28758<br>[0;107494] | 25441<br>[0;86484] |
| $\hat{T}_{AM}$ | 64496<br>[5749;141447] | 67230<br>[6518;141884] | 70088<br>[5740;143504] | 61231<br>[7853;136819] |
| $\hat{T}_{split}$ | 83594<br>[18549;143948] | 88566<br>[15300;146844] | 94526<br>[16459;147573] | 79413<br>[16947;142844] |
| $\hat{Q}$ | 0.049<br>[0;0.175] | 0.048<br>[0;0.180] | 0.053<br>[0;0.178] | 0.037<br>[0;0.166] |

*Table S6: Weight of each demographic model in posteriors for each pair of crow species from Vijay et al (2016) and Poelstra et al (2014) (=Poelstra comp).*

| Demographic model weight | AM | IM | SC | SI |
|---|---|---|---|---|
| RX | 0.04132532 | 0.49096738 | 0.44987948 | 0.01782782 |
| OP | 0.09784093 | 0.47982347 | 0.38247273 | 0.03986287 |
| XO | 0.09507434 | 0.45523509 | 0.41617606 | 0.03351451 |
| Poelstra comp | 0.07057804 | 0.41757511 | 0.48631229 | 0.02553457 |
